# Feedback Inhibition Underlies New Computational Functions of Cerebellar Interneurons

**DOI:** 10.1101/2022.03.03.482855

**Authors:** Hunter E. Halverson, Jinsook Kim, Andrei Khilkevich, Michael D. Mauk, George J. Augustine

## Abstract

The function of a feedback inhibitory circuit between cerebellar Purkinje cells and molecular layer interneurons (MLIs) was defined by combining optogenetics, neuronal activity recordings both in cerebellar slices and *in vivo*, and computational modeling. Purkinje cells inhibit a subset of MLIs in the inner third of the molecular layer. This inhibition is non-reciprocal, short-range (less than 200 μm) and is based on convergence of 1-2 Purkinje cells onto MLIs. During learning-related eyelid movements *in vivo*, the activity of a subset of MLIs progressively increases at the same time that Purkinje cell activity decreases, with Purkinje cells usually leading the MLIs. Computer simulations indicate that these relationships are best explained by the feedback circuit from Purkinje cells to MLIs and that this feedback circuit plays a central role in making cerebellar learning efficient.

---

Local inhibitory interneurons are ubiquitous throughout the brain and play diverse and fundamental roles in neuronal information processing^1^. Within the cerebellar cortex, Golgi, basket and stellate cells have been identified as inhibitory interneurons. While Golgi cells are found in the granule cell layer, basket and stellate cells reside within the molecular layer of the cerebellar cortex. These molecular layer interneurons (MLIs) form inhibitory synapses with each other, as well as with Purkinje cells (PCs), the only output cells of the cerebellar cortex. Whereas specific computational roles have been proposed for Golgi cells, including control of granule cell threshold and temporal coding^2, 3^, proposals regarding the functions of MLIs generally have been limited to the relatively simplistic notion that they contribute to feed- forward inhibition of PCs. Moreover, the function of stellate and basket cells generally are not considered separately in theories of cerebellar cortex.

Following up on suggestions that MLIs may receive inhibitory input from PCs^4, 5^, we present evidence from high-speed optogenetic circuit mapping and paired recordings in cerebellar slices that some MLIs in the inner third of the molecular layer, but not stellate cells in the outer molecular layer, are inhibited by feedback from PCs. Because these MLIs, in turn, inhibit other PCs, our results suggest that PCs influence each other through disinhibition of MLIs. Consistent with this disinhibitory circuit, *in vivo* a subset of MLIs fire strongly anti-phasically with, and usually after, PCs during eyelid conditioning. Finally, large-scale computer simulations predict that PC feedback inhibition of MLIs enhances the ability of the cerebellum to learn and to facilitate more efficient information storage.

## RESULTS

### Photostimulation of Purkinje cells

We used mice that express the light-activated cation channel, ChR2, exclusively in PCs to enable selective photostimulation of PCs *in vitro*. These mice were obtained by crossing transgenic mice expressing Cre recombinase behind a PC-specific promoter (PCP2)^6^ with another transgenic mouse line expressing ChR2-H134R behind a floxed stop cassette^7^. Similar results were obtained using a second PCP2 -cre line^5^. In these mice, light pulses generate photocurrents sufficient to evoke action potentials in PCs^8^. We examined ChR2 expression by imaging the yellow fluorescent protein fused to the ChR2; virtually all PCs strongly expressed ChR2 in their somata and dendrites, with no fluorescence evident in MLI somata within the molecular layer (**Fig. 1a**). Whole-cell patch clamp recordings in cerebellar slices from these double transgenic mice indicated that photostimulation evoked action potentials in every PC examined (20/20), but was incapable of evoking action potentials in MLI (0/406).

**Figure 1.**
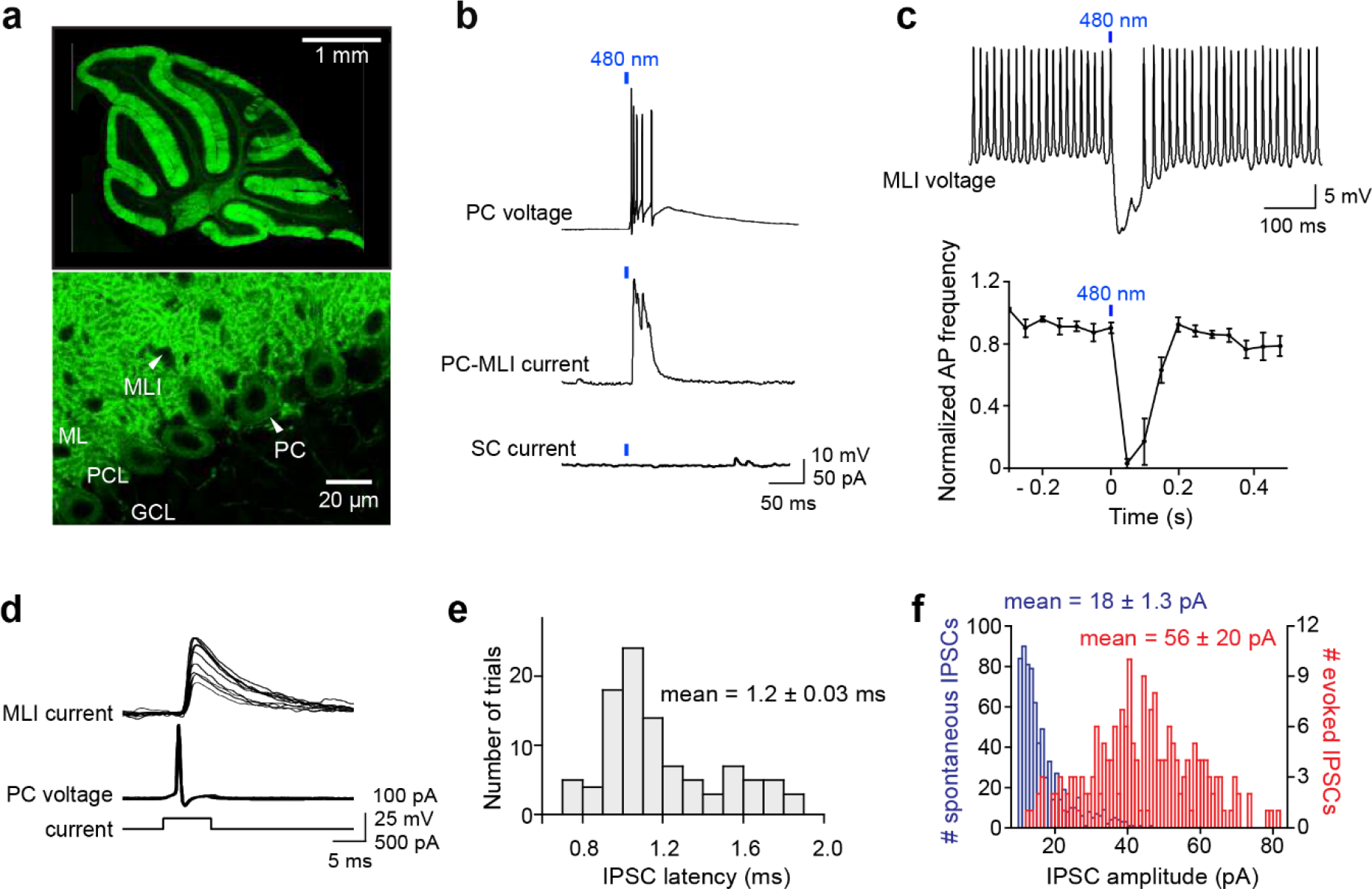
Optogenetic interrogation of Purkinje cell to MLI circuit. (a) Selective expression of ChR2 in cerebellar PCs. Top: Image of ChR2-YFP fluorescence (green) in a sagittal cerebellum section from PCP2-Cre; Ai32 double transgenic mice. Strong ChR2 expression was observed throughout entire cerebellar cortex, especially in the molecular layer where PC dendrites are located. Bottom: higher-magnification image shows expression of ChR2-YFP in their soma (arrowhead) and dendrites. Small black holes (arrowhead) in the molecular layer represent somata of MLI that do not express ChR2-YFP. ML, molecular layer; PCL, Purkinje cell layer; GCL, granule cell layer. (b) Brief illumination (480 nm, 5 ms, 9.9 mW/mm^2^) evoked action potentials in ChR2- expressing PCs (top). Photostimulated PCs induced IPSCs in PC-MLI cells (center) but not in stellate cells (bottom). (c) Top: Photostimulation of PCs (at blue bar) inhibited firing in a postsynaptic MLI. PC-MLI was depolarized by a depolarizing current (20 pA) to sustain action potential firing. Bottom: Activation of PCs input (at blue bar) was sufficient to briefly but completely inhibit action potential firing in postsynaptic PC-MLI cells. Points indicate means and error bars represent SEMs (n = 4). (d) Superimposed traces of recordings from a connected PC and PC-MLI pair. Action potentials in a presynaptic PC (middle traces), caused by depolarizing current pulses (bottom traces), induced IPSCs in the postsynaptic PC-MLI (top traces). PC-MLI holding potential was -50 mV. (e) Distribution of IPSC latencies measured in 7 PC-MLI pairs (140 trials). Mean IPSC latency was 1.2 ms with relatively low synaptic jitter (0.03 ms). (f) Distribution of amplitudes of spontaneous (blue) and evoked (red) IPSCs in PC-MLIs.

We characterized synaptic connectivity between PCs and MLIs in sagittal slices of the cerebellar vermis, based on previous anatomical studies indicating that PC axon collaterals are oriented in the sagittal plane^5, 9^. We started by using relatively large spots of blue light (∼0.2 mm^2^ area) to photostimulate a large number of PCs (**Fig. 1b**, top), while measuring postsynaptic responses in MLIs. Inhibitory postsynaptic currents (IPSCs) were observed only in MLIs located in the inner one-third of the molecular layer, where basket cells are located (**Fig. 1b**, center). IPSCs were abolished by GABAA receptor blockers, such as bicuculline and gabazine, but were unaffected by the glutamate receptor blocker kynurenic acid, indicating that the IPSCs are monosynaptic and mediated by GABAA receptors. Connectivity between PCs and MLIs was observed in 16% of the 371 MLIs examined in the inner third of the molecular layer, where basket cells are located, in mice aged 3 weeks or older. No connections were detected (n = 35) between PCs and MLIs located in the outer two-thirds of the molecular layer, where stellate cells are located (**Fig. 1b**, bottom). We refer to the subset of MLIs receiving PC inputs as “PC-MLIs” (**Fig. 1b**, middle). The somata of PC-MLI were 60 ± 2.9 μm (mean ± SEM, n = 59) above the PC layer and thus are not candelabrum cells, interneurons whose somata reside within the PC layer^10^. Inhibitory feedback from PCs to PC- MLIs was also observed in the lateral cerebellum: a total of 8 connections were detected among 34 recordings in the inner molecular layer of lobulus simplex, crus I, and crus II lobules. This 24% rate of connections compares favorably to the ∼16% observed in 371 recordings in the vermis. Therefore, the PC-to-PC-MLI feedback circuit seems widely distributed throughout the cerebellum and may play a general role in cerebellar computation. To determine how feedback from PCs affects the activity of PC-MLI, we photostimulated PCs with brief light pulses (5 ms) while depolarizing PC-MLIs to evoke rapid action potential firing. Light-evoked synaptic input from PCs was capable of completely suppressing PC-MLI activity (**Fig. 1c**).

We then used recordings from pairs of connected PCs and MLIs for detailed characterization of the inhibitory feedback circuit. Although recordings from connected pairs were rarely obtained (see below), seven successful experiments (6 whole-cell recordings and 1 cell-attached recording from PCs) allowed us to directly define the functional attributes of this circuit. Single action potentials (simple spikes) evoked in presynaptic PCs induced IPSCs in postsynaptic MLIs (**Fig. 1d**). IPSC latency was brief, with a mean of 1.15 ± 0.03 ms (**Fig. 1e**). IPSCs were quite variable in amplitude (**Fig. 1f**, red), with a coefficient of variation (CV) of 0.39 ± 0.08 and peak amplitude of 56.1 ± 20.3 pA at -50 mV (n = 7). This indicates a synaptic conductance of 2.13 ± 0.61 nS. The mean amplitude of spontaneous (presumed miniature) IPSCs was 18.0 ± 1.3 pA (**Fig. 1e**, blue), indicating a quantal content (ratio of amplitudes of evoked/spontaneous IPSCs) of 3.0 ± 0.9 for evoked IPSCs. Recalculating based on IPSC integral, rather than peak amplitude, raised the estimate to 6.7; this reveals a substantial contribution of asynchronous GABA release^11^. These unitary synaptic properties are similar to those reported for synapses formed between PCs by PC axon collaterals^5, 9, 12^. However, unlike PC-PC synapses^5, 9^, the PC-to-PC-MLI synapse exhibited very few failures (**Fig. 1d**).

This is a consequence of the relatively high quantal content and large readily releasable pool (RRP) of synaptic vesicles (99 ± 16 quanta; n = 15; Fig. 2h), as well as a relatively low release probability (0.07), determined from the ratio of quantal content to RRP size.

**Figure 2.**
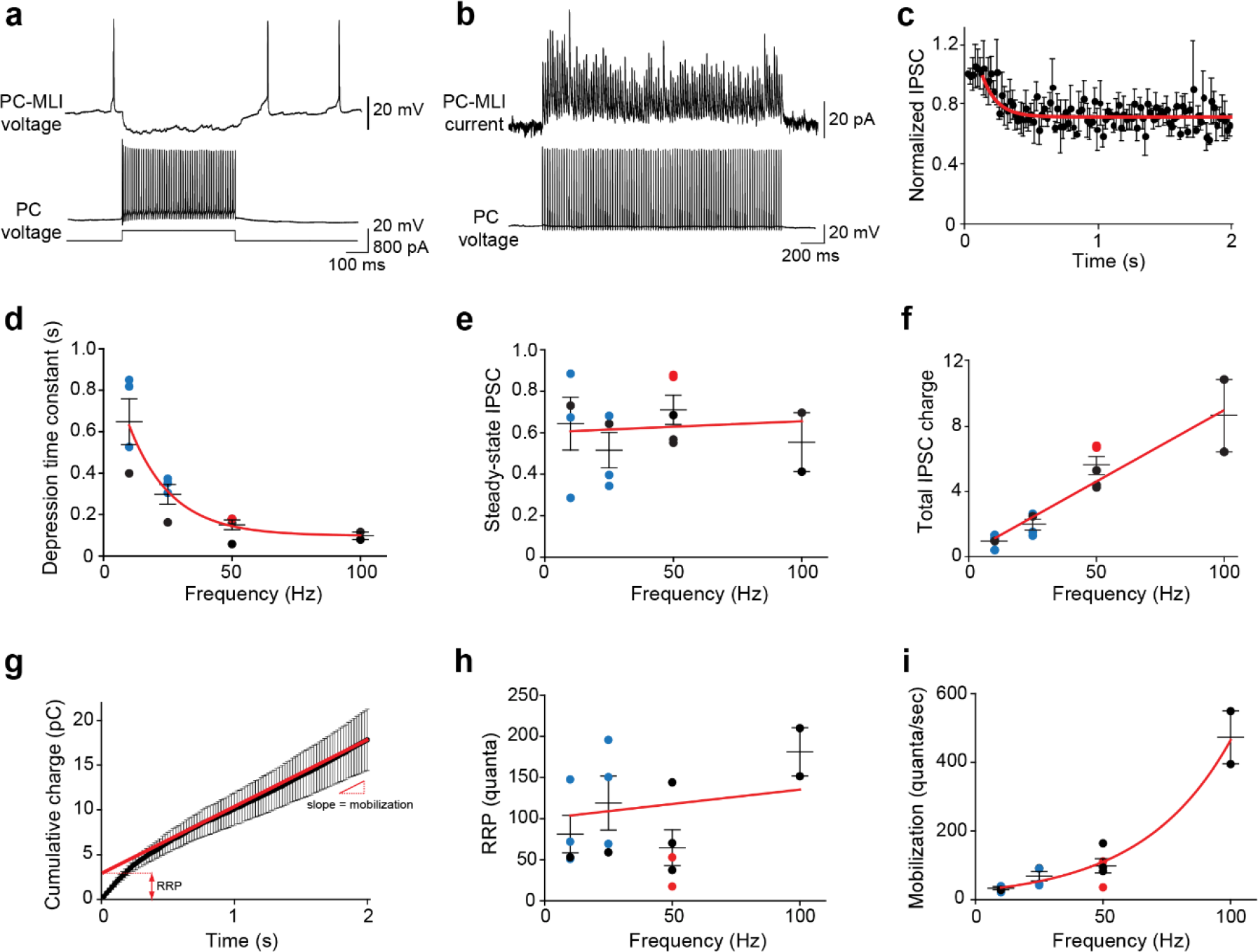
Frequency-independent depression at Purkinje cell-to-PC-MLI synapses. (a) Example of recordings of synaptic transmission at a connected PC and PC-MLI pair. 100 Hz firing of the presynaptic PC for 500 ms produced a sustained inhibition of the postsynaptic PC-MLI. (b) Average IPSCs (10 trials) evoked by 100 stimuli (50 Hz for 2 sec) from a connected PC and PC-MLI pair. During such trains of synaptic activity IPSCs often initially showed facilitation, indicated by an increase in IPSC amplitude, which was followed by relatively mild synaptic depression, indicated by a reduction in IPSC amplitude. (c) Kinetics of synaptic depression during sustained synaptic activity. IPSC amplitudes were normalized to the first IPSC in each stimulus (IPSCn /IPSC1) during 50 Hz stimulation. Points indicate mean values and bars are ± SEMs (n = 5), while red line is an exponential fit to the time course of depression of IPSC amplitude. (d) Depression time constants were measured from the exponential fit, as shown in (c), at different stimulation frequencies. Symbols represent data collected via different methods of stimulation of presynaptic PCs: blue - photostimulation; red - current injection in dual recordings; black - extracellular stimulation. Red line is an exponential fit to the data. (e) Steady-state amplitudes of IPSCs measured during stimulus trains of different frequencies; values are normalized to the first response in a train, as in (c). Red line indicates a linear fit to the data; slope of nearly zero indicates no significant effect of the frequency of synaptic activity on steady-state IPSC amplitudes. (f) Steady-state charge transfer (IPSC amplitude x frequency) measured at different frequencies of synaptic activity. Colored symbols represent mean values determined for individual experiments, using the color code described for (d), and bars indicate ± SEM. Red line represents a linear fit to the data and indicates a dramatic effect of the frequency- dependent enhancement of cumulative synaptic transmission. (g) Kinetics of synaptic charge transfer during a stimulus train. Points indicate IPSC charge integrated over time during 50 Hz stimulation, while bars indicate ± SEM (n = 5). The last 20 data points were fit by linear regression (red line) to estimate the size of the readily releasable pool (RRP; y-intercept) and mobilization rate (slope). (h) RRP size were measured, using the analysis shown in (g), at different stimulus frequencies. Red line indicates a linear fit to the data; slope of approximately zero indicates no significant effect of the frequency of synaptic activity on RRP size. (i) Rate of synaptic vesicle mobilization from the reserve pool to the RRP was estimated using the analysis shown in (g). Points represent mean values determined for individual experiments, while bars are ± SEM. Red line is an exponential fit to the data and indicates a large effect of the frequency of synaptic activity on the rate of vesicle mobilization.

Because PCs fire repetitively at rates up to 100 Hz *in vivo*^13, 14^, we determined how sustained activity affected transmission at the PC-to-PC-MLI synapse at near-physiological temperature (34–35°C). Trains of presynaptic PC activity (10-100 Hz) initially caused synaptic facilitation, which was followed by a mild depression of IPSC amplitude (Fig. 2a,b). Depression followed an exponential time course, with the rate of depression depending upon stimulus frequency (Fig. 2c,d). Remarkably, IPSC amplitude was sustained at approximately half of its initial strength even during prolonged activity, independent of stimulus frequency (Fig. 2e). As a result, the steady-state amount of synaptic transmission – measured as total IPSC charge - paradoxically increased at higher frequencies (Fig. 2f). By measuring cumulative synaptic charge during repetitive activity (Fig. 2g), we determined the size of the RRP (Fig. 2h) and the rate of mobilization of synaptic vesicles from a reserve pool (Fig. 2i). This analysis revealed that PCs sustain their transmission via an activity-dependent acceleration of synaptic vesicle mobilization from the reserve pool to the RRP, similar to synapses between PCs and deep cerebellar nuclear neurons^15^. Thus, PCs can inhibit PC-MLIs even during high-frequency activity, thereby maintaining an inverse relationship between the activities of these two neuron types.

### Mapping the spatial organization of the Purkinje cell-MLI circuit

The circuit between presynaptic PCs and postsynaptic MLIs was mapped by scanning small spots of laser light (405 nm, 4 ms duration, ∼1 μm diameter in the focal plane) to evoke action potentials in presynaptic PCs expressing ChR2^16, 17^. Under these conditions, 1.2 µW or more of laser power could reliably evoke action potentials in PCs in cerebellar slices from mice 3 weeks or older. Optimal spatially-resolved photostimulation^17^ occurred at a laser power of 3 µW: at this laser power, light spots were capable of evoking action potentials at any location on the dendrite or cell body of individual PCs (**Fig. 3a**, position 2), aside from their axons. In contrast, positioning the light spot more than 30 μm away from the PC soma along the PC layer failed to evoke action potentials (**Fig. 3a**, positions 1 or 3).

**Figure 3.**
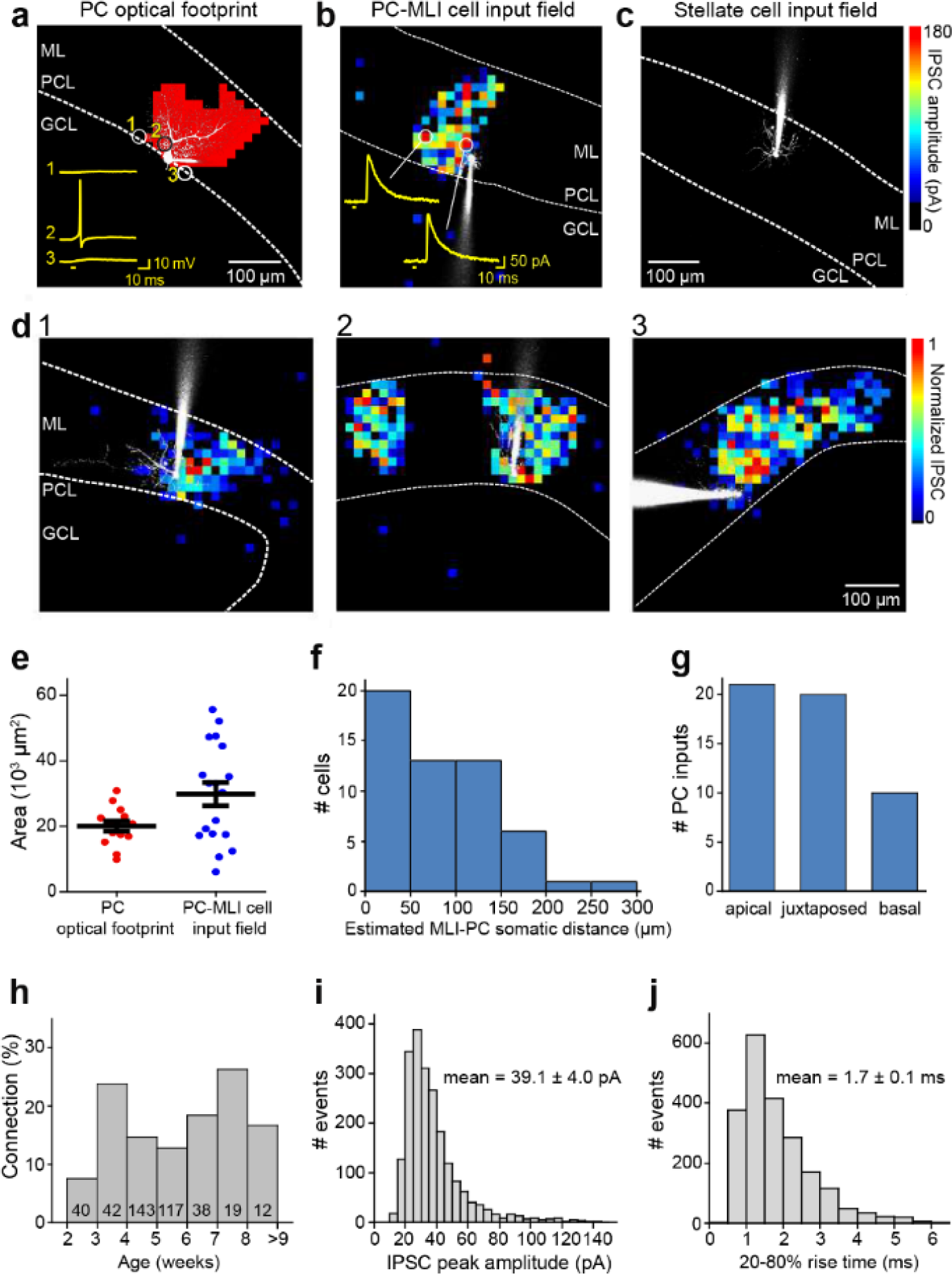
Spatial organization of Purkinje cell to MLI circuit. (a) Scanning a brief laser light spot (405 nm, 3 µW, 4 ms duration) across a cerebellar slice revealed locations (red pixels) where photostimulation evoked action potentials in a ChR2- expressing, dye-filled PC (white). Numbered traces (yellow) in the inset show membrane potential changes produced when laser spot was located at the indicated pixels. Bar below traces indicates time of light flashes. Action potentials were evoked in the cell body and entire dendritic region of the PC. ML, molecular layer; PCL, Purkinje cell layer; GCL, granule cell layer. (b) Inhibitory input map in a postsynaptic PC-MLI was created by correlating light spot location with amplitude of IPSCs evoked by focal photostimulation of PCs. IPSC amplitudes are encoded in the pseudocolor scale shown at right. Traces show IPSCs evoked at the indicated locations and bars indicates time of light flashes. (c) No inhibitory input from PCs was detected in stellate cells. (d) 1-3: Varied spatial organization of PC inputs to PC-MLIs. Maps illustrate IPSC amplitudes (pseudocolor scale at right) evoked in three different dye-filled PC-MLIs (white). (e) Comparison of input field areas of PC-MLIs (blue, n = 18) with PC optical footprints (red, n = 14). (f) Distance between postsynaptic PC-MLIs and presynaptic PCs. PC soma location was estimated from the center of the input field, projected down to the Purkinje cell layer. (g) Orientation of postsynaptic PC-MLIs, relative to presynaptic PCs. (h) Connectivity between PCs and PC-MLIs measured at different ages. Numbers inside bars indicate sample sizes. (i) Distribution of amplitudes of IPSCs evoked by focal photostimulation of PCs. Holding potential was -40 mV. (j) Distribution of rise times of light-evoked IPSCs.

The spatial organization of the circuit was mapped by scanning the laser light spot to evoke action potentials in presynaptic PCs, while recording IPSCs in MLIs. A light-evoked IPSC indicated a synaptic connection between a PC photostimulated at that location and postsynaptic PC-MLI; thus, correlating light spot location with the amplitude of IPSCs mapped the position of presynaptic PCs^16, 17^ (**Fig. 3b**). Recordings from stellate cells in the outer molecular layer yielded blank maps (**Fig. 3c**), again indicating an absence of connections between PCs and stellate cells (n = 35). IPSCs were evoked in PC-MLI only when the light spot was positioned within the molecular and PC layers, where PCs are located. Presynaptic PC inputs to PC-MLI were most often revealed (59%) as a single cluster of pixels similar in size to the light-sensitive area of a single PC (**Fig. 3d1**), occasionally (12%) as two discrete input areas (**Fig. 3d2**) or a relatively large contiguous input area (29%; **Fig. 3d3**). Within an input field, there were no obvious “hot spots” of synaptic input, consistent with the homogeneous light sensitivity of individual presynaptic PCs (**Fig. 3a**).

To estimate the number of PC inputs converging on a PC-MLI, we compared the area of the synaptic input field (**Fig. 3d**) with the area over which light spots could evoke action potentials in single PCs (optical footprint^17^; **Fig. 3a**). Mean input field area was 1.5 times larger than the mean area of PC optical footprints, suggesting that on average a PC-MLI receives inhibitory input from only one or two converging PCs (**Fig. 3e**). This is consistent with the observed spatial organization of input maps shown in **Fig. 3d**. The spatial organization of the PC-to-PC-MLI circuit is different from that of the MLI-to-PC circuit^17^: presynaptic PCs were largely found within 200 µm of their postsynaptic PC-MLI (**Fig. 3f**) and were asymmetrically distributed, with most presynaptic inputs originating from either juxtaposed or apically located PCs (**Fig. 3g**). These observations are consistent with the anatomy of PC axon collaterals^5, 9^.

Synaptic inputs from PCs to PC-MLI could be detected as early as postnatal day 14 (P14; **Fig. 3h**), a time when MLIs are still developing^18, 19^. However, the rate of connectivity at ages earlier than P21 (7.5%) was somewhat lower than at later ages (16%), suggesting that the circuit is established during the time that MLIs mature. After P21, the probability of detecting functional connections between PCs and PC-MLI was relatively constant up to P77 (**Fig. 3h**), well after MLIs have fully matured^18, 19^. Thus, this synaptic circuit is functional into adulthood. No feedback inhibitory connections were observed between PCs and stellate cells at any time between postnatal weeks 3 to 9.

The large sample of IPSCs collected during our mapping experiments allowed further characterization of synaptic transmission at the PC-to-MLI synapse. Within the PC input fields, IPSC amplitude varied substantially from pixel to pixel but this variability was not due to variations in photostimulation of presynaptic PCs (**Fig. 3a**). The CV of light-evoked IPSCs was 0.28 ± 0.02 in 18 experiments; this is similar to the CV of IPSCs observed in our paired recordings from PCs and PC-MLIs (**Fig. 1d**) and provides a second indication of highly variable transmission at the PC-to-MLI synapse. Mean IPSC amplitude, measured at a holding potential of -40 mV, was 39.1 ± 4.0 pA (**Fig. 3i**; 2128 IPSCs in 18 recordings). This corresponds to a synaptic conductance of 1.15 ± 0.12 nS, which is not significantly different from that measured in paired recordings (p = 0.17, Welch’s t-test). IPSC rise time (20-80%) was relatively fast (1.69 ± 0.13 ms, **Fig. 3j**), similar to IPSCs produced at PC-to-PC synapses^5, 9, 12^ but more rapid than the MLI-to-PC connection (5.6 ± 0.5 ms time-to-peak; see Fig. 5c of ref. 17).

### Non-reciprocal connections between Purkinje cells and MLIs

Given that MLIs provide feedforward inhibition to PCs, we next asked whether circuits between PCs and MLIs are reciprocal, i.e. whether PCs innervated by a given MLI provide feedback inhibition to the same MLI. For this purpose, we combined paired electrophysiological recordings with optogenetic mapping. Due to the abundance of PCs, we used mapping to increase the probability of detecting functional PC-MLI connections: after finding a PC-MLI and identifying the location of its presynaptic PC by scanning the photostimulating laser spot, as in **Fig. 3b,d**, recordings were made from the somata of PCs within the area where the light spot evoked IPSCs. Among 139 paired recordings made via this strategy, synaptic inputs from PCs to MLIs were detected in 6 cell pairs (excluding one cell-attached PC recording). This 5% success rate is an underestimate of PC-to-PC-MLI connectivity; although our optogenetic mapping identified the volume in which presynaptic PCs were located, there was still a low likelihood of sampling the 1 or 2 presynaptic PCs within the dozens of PCs within this volume. The results from one successful experiment are shown in **Fig. 4a**. Action potentials evoked in the PC by depolarizing current pulses elicited IPSCs in the postsynaptic PC-MLI (**Fig. 4a**, upper traces). However, evoking action potentials in the PC-MLI did not elicit IPSCs in the PC, indicating that this pair did not have a reciprocal connection (**Fig. 4a**, lower traces). This was the case for all 6 coupled PC/PC-MLI pairs.

**Figure 4.**
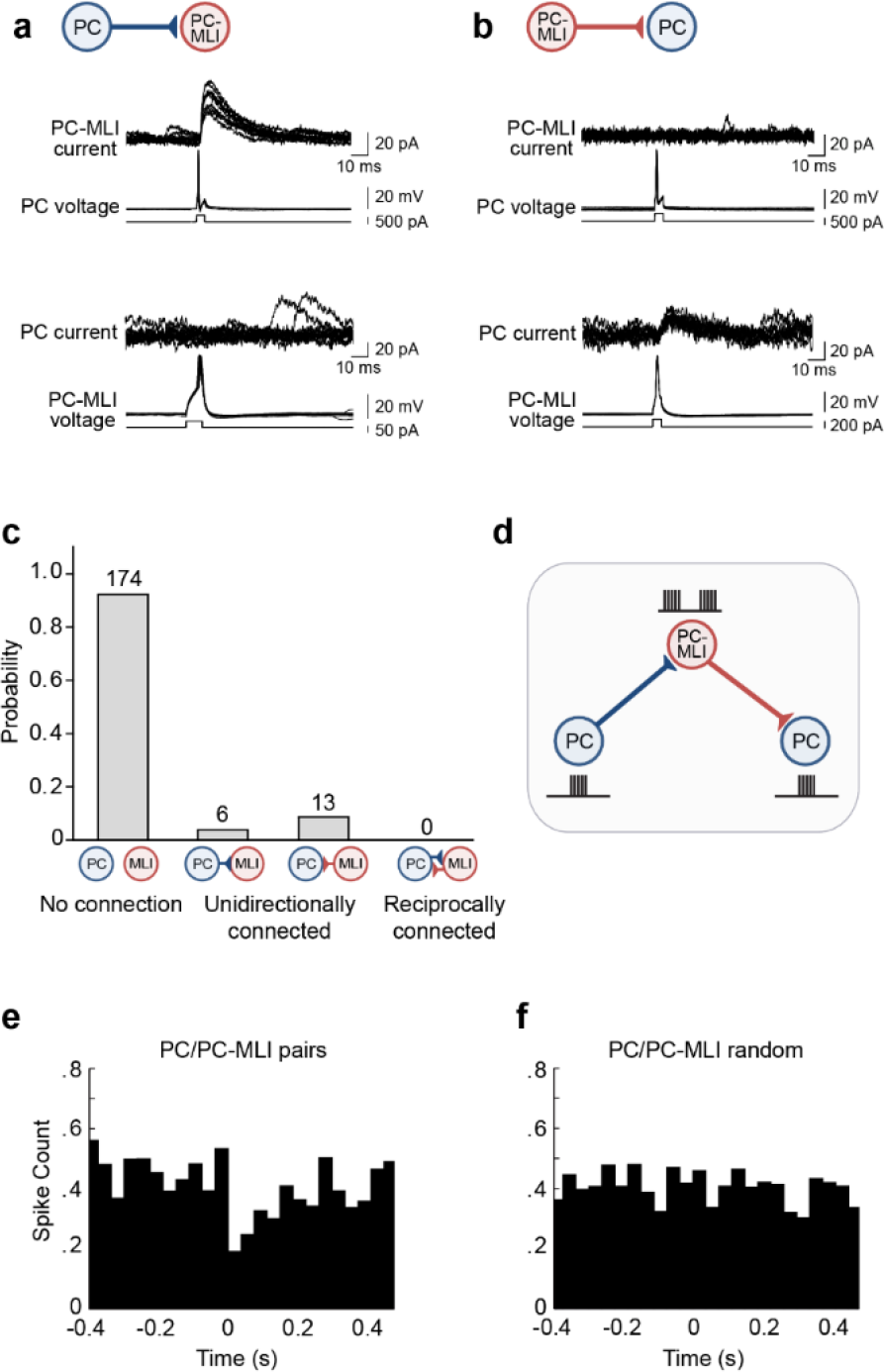
Evidence for non-reciprocal arrangement of inhibitory circuits between MLIs and Purkinje cells. (a) Action potentials evoked in a presynaptic PC induced IPSCs in a postsynaptic PC-MLI (top). However, action potentials in the PC-MLI did not evoke IPSCs in the PC (bottom). (b) In random paired recordings, inhibitory inputs from PC-MLIs to PCs could be observed (bottom). Again, these were non-reciprocal because stimulating the PC did not evoke IPSCs in the PC-MLI (top). (c) Probability of connections between PCs and MLIs observed in dual recordings. Only unidirectional connections were detected (n = 193). (d) Proposed organization of circuits between PCs and PC-MLIs. PC-mediated inhibition of PC-MLI activity may disinhibit neighboring PCs. (**e**, **f**) Temporal relationship between action potential firing in connected PC/PC-MLI pairs in brain slices. Cross-correlations (at time lag = 0) between presynaptic PC and postsynaptic PC-MLI pairs (e) and activity in the same pair of cells after randomly shuffling the timing of PC spikes (f). The inhibitory connection between PCs and PC-MLIs yielded a strong, but transient, inverse activity correlation.

As an alternative strategy, we randomly selected PCs within 100 µm of an MLI located in the inner third of the molecular layer. Synaptic inputs from MLIs to PCs could be detected, evident as IPSCs evoked in PCs in response to MLI stimulation (**Fig. 4b**, lower traces, 13/54 pairs). However, in such cases, action potentials in the postsynaptic PC invariably failed to evoke reciprocal IPSCs in the MLI (**Fig. 4b**, upper traces). In 13 experiments where stimulation of a MLI in the inner 1/3 of the molecular layer evoked IPSCs in a PC, in no case did stimulation of the PC produce IPSCs in the MLI partner (**Fig. 4c**). Overall, in a total of 193 paired recordings, only unidirectional connections (either PC-to-MLI or MLI-to-PC) or unconnected pairs were detected; in no case were reciprocally connected PCs and MLIs observed (**Fig. 4c**). Although limited by the number of recordings from connected pairs (19), our results suggest that the inhibitory circuits between PCs and MLIs are non-reciprocal, yielding a connectivity where PCs can influence other PCs through disinhibition, while not influencing their own behavior through recurrent inhibition (**Fig. 4d**).

To understand how PC activity influences that of postsynaptic PC-MLIs, we examined the temporal relationship between spikes in PCs and in PC-MLIs (4 cell pairs). This cross-correlation analysis revealed that inhibitory inputs from PCs to PC-MLI cells produced a clear inverse correlation in the activity of these two cell types (**Fig. 4e**). Shuffling the timing of PC spikes eliminated this effect (**Fig. 4f**), indicating that the inverse correlation normally observed after a PC action potential is genuine and is not an artifact of our analysis method.

### Cerebellar processing suggested by circuit simulations

The connectivity revealed by the slice experiments suggests that PC-MLIs receiving feedback inhibition from a nearby PC should fire out of phase with PCs, increasing their activity when PC activity decreases. This anti-correlated pattern of activity between PCs and PC-MLIs could have important implications for cerebellar information processing. To determine the function of this feedback circuit, we first used a large-scale computer simulation of cerebellar circuitry (**Fig 5a**) to predict neuronal activity during eyelid conditioning, a well-characterized cerebellar-dependent form of motor learning. The simulation has been employed over many years and has been optimized to successfully model many aspects of cerebellar processing, including bidirectional learning, adaptive timing and roles for multiple sites of plasticity, short-term plasticity and recurrent feedback^22–28^. To this simulation we added feedback from PCs to PC- MLIs. Based on the location of PC-MLIs within the inner one-third of the molecular layer, in the simulations the connectivity of PC-MLIs was assumed to be identical to that of basket cells. Simulated mossy fiber and climbing fiber inputs were based on responses of these inputs measured during eyelid conditioning in rabbits. These simulations also included excitatory recurrent collaterals from the deep cerebellar nuclei, which deliver to the cerebellar cortex - as mossy fiber input - a copy of output that can be used for sequence learning^29, 30^. These collaterals were included in the same proportion as mossy fibers representing the conditioned stimulus (CS): 5.2% of all mossy fibers^31^ (see Supplementary Text for further details).

**Figure 5.**
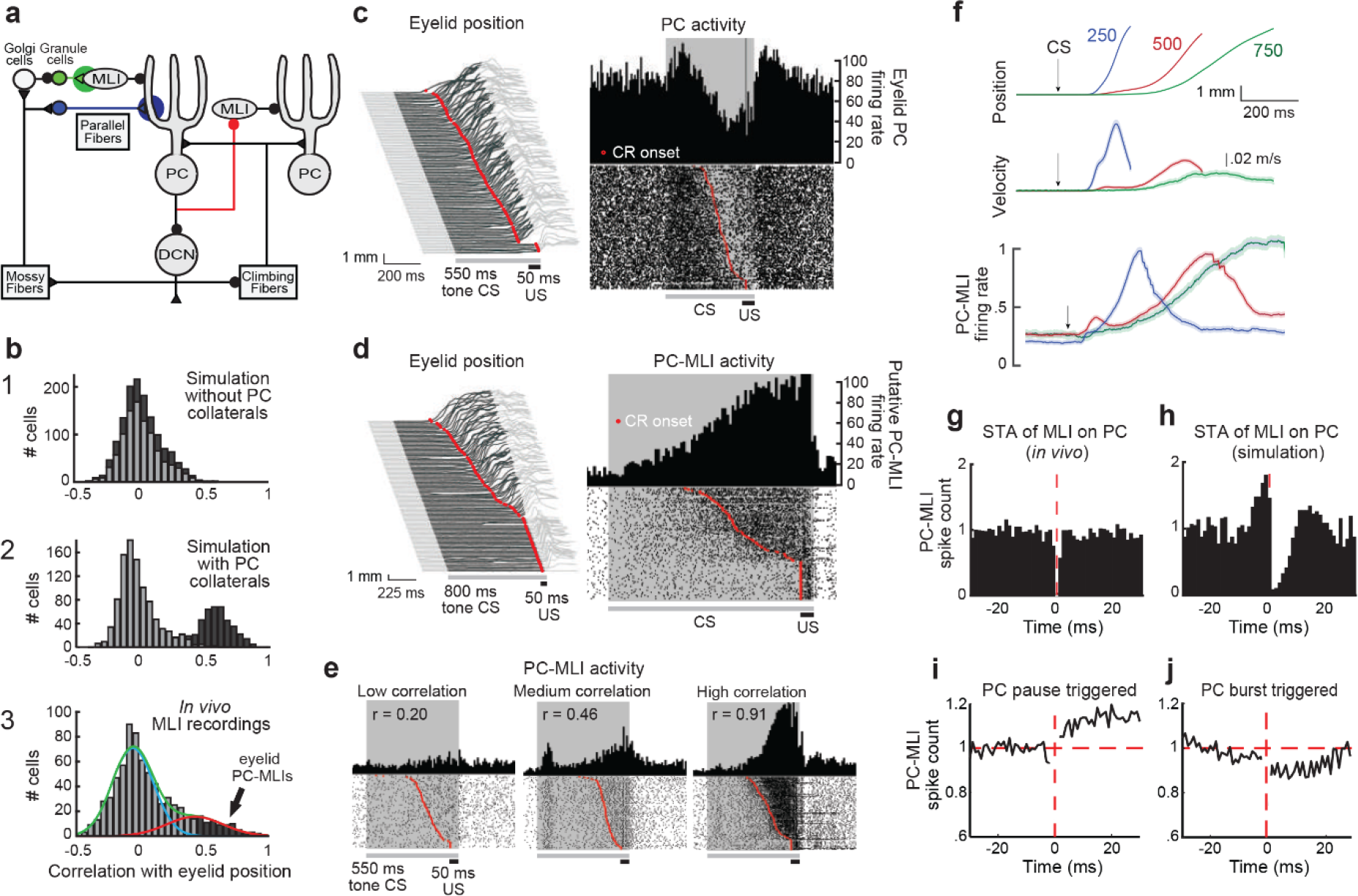
Molecular layer interneuron *in vivo* recordings and the relationship with behavior. (a) Circuit diagram showing main connections and sites of plasticity for the computer simulation of the cerebellum. Green indicates potential plasticity at granule to MLI synapses and blue indicates plasticity at granule cell to PC synapses. Red indicates the inhibitory feedback connection from PCs to MLIs. (b) Cross-correlations between MLI activity and eyelid CRs. The top two panels show data from a computer simulation of the cerebellum that differed only in that one simulation lacked inhibitory feedback from PCs to MLIs (1) and the other had feedback from PCs to MLIs (2). In well-trained simulations, correlations were calculated between MLI activity and the predicted CRs. Simulations with the PC to MLI feedback showed two distributions, one from a subset of MLIs that is shifted in the positive direction (dark gray, middle panel) and the other centered around zero (light gray). The simulation without feedback showed one distribution centered around zero for all MLIs (light and dark gray). (3) Distribution of MLI correlations recorded *in vivo* for all ISIs can be fit with two distributions, one shifted in the positive direction (red curve) and one centered around zero (blue curve) similar to the simulation with the PC to MLI feedback. The cutoff for categorizing a MLI as a PC-MLI was r ≥ .43, where there was a 95% chance that the two fitted distributions were different. (c) Left – Eyelid movements (vertical) in response to conditioned stimuli (CS) and unconditioned stimuli (US), sorted by the latency of the conditioned response (CR; red dots). Upward deflection indicates eyelid closure, with closure indicated by dark gray reflecting CRs. Right - Peri-stimulus time histogram (upper; 10 ms bins) and raster plot (lower) of an eyelid PC recording from all trials of the eyelid conditioning session shown at left. For this and all subsequent raster plots, the gray bar under the plot indicates duration of the tone stimulus and the black bar indicates the duration of the eyelid stimulation. (d) Peri-stimulus time histogram (10 ms bins) and raster plot of a PC-MLI recording (right), along with the behavior from all trials of the eyelid conditioning session during the recording (left), sorted by the latency of the conditioned responses (red dots). (e) Peri-stimulus time histograms (10 ms bins) and raster plots sorted by response latency (red dots) showing three example MLI recordings. The first example (left; r = 0.20) is from the low-correlation MLI distribution from the in vivo recordings. The last two examples (r = 0.46 and 0.91) were operationally defined as eyelid PC-MLIs due to their high correlation with behavior. (f) Average eyelid position (top), velocity (middle) and normalized average eyelid PC-MLI activity (bottom) shown for 3 different ISIs. The shaded region represents the 95% confidence intervals and arrows represent time of CS onset. (**g** and **h**) Average cross-correlograms calculated from significantly modulated pairs of MLIs and eyelid PCs. Background activity of putative PC-MLI is triggered off PC simple spikes (vertical red line), for PC-MLI and PC pairs recorded on the same tetrode (g) and from the computer simulation (h). Decrease in MLI activity after a PC simple spike indicates an inverse relationship between the activity of the two cell types. (**i and j**) Average PC pause-triggered (i) and burst-triggered (j) cross-correlograms of putative PC-MLI background activity. Plots show average PC-MLI activity across significant pairs recorded on the same tetrode. Pauses in PC simple spikes resulted in an increase in mean PC-MLI activity, while bursts of PC simple spikes resulted in a decrease in mean PC-MLI activity.

To examine the role of PC-to-MLI feedback, we next tested simulations with and without feedback inhibition between PCs and PC-MLIs (red in **Fig. 5a**), using a variety of inter- stimulus intervals (ISIs) for training to yield CRs that varied widely in their timing. Without PC- MLI feedback inhibition, the activity of most MLIs tended to be uncorrelated with CRs (**Fig. 5b1**). On the other hand, simulations including PC feedback inhibition of PC-MLIs yielded a bimodal distribution of correlations, with a subpopulation of MLIs having a high correlation with eyelid CRs (mean r = 0.60 ± 0.06 SEM) and other MLIs having minimal correlation with CRs (mean r = -0.02 ± 0.01 SEM) (**Fig. 5b2**). Therefore, MLIs with activity correlated with CRs are likely to be PC-MLIs whose activity increases during eyelid conditioning because presynaptic eyelid PCs become less active. The simulations further indicate that this inverse relationship arises from the PC-to-PC-MLI feedback circuit, rather than from feedforward inhibition of PCs by MLIs. Finally, this result predicts that decreases in PC activity should precede increases in

MLI activity during expression of learned responses. These predictions are experimentally testable and provide a basis for interpreting the *in vivo* recordings of neuronal activity during delay eyelid conditioning in rabbits described in the next section.

### Bimodal responses of MLIs *in vivo*

We used *in vivo* recordings of cerebellar neuron activity after eyelid conditioning to test predictions of our model based on the PC-to-PC-MLI feedback circuit. **Fig. 5c** (left) illustrates eyelid responses after conditioning. In this and subsequent figures, the gray shaded area represents presentation of the CS (tone) and responses are sorted according to the onset of conditioned closure of the eyelid (eyelid CR), indicated by the upward deflections (dark gray) during the CS. Onset of eyelid movement is indicated by red dots. A peristimulus time histogram (**Fig. 5c**, upper right) illustrates an eyelid PC whose activity decreased during the CR and a raster plot (**Fig. 5c**, lower right) reveals that these changes in PC activity were highly correlated, on a trial-by-trial basis, with the time of onset of eyelid movement.

Because eyelid CRs are driven by these learning-dependent decreases in eyelid PC activity^32, 35^, PC-MLIs should fire inversely with eyelid PCs and this activity should be highly correlated with eyelid CRs. We recorded MLI responses during learned expression of eyelid CRs to test the prediction of our simulations that a fraction of MLIs respond in ways consistent with PC-to-PC-MLI feedback. In extracellular recordings, MLI activity can be distinguished by an established algorithm that statistically categorizes neurons according to their baseline firing properties^36^: MLIs are identified by their relatively irregular firing. Indeed, we observed a subset of MLIs whose activity increased in high temporal correlation with CR expression. **Fig. 5d** shows an example of such a cell, evident from the peristimulus time histogram (upper right) and raster plots (lower right) of action potential firing, and the corresponding eyelid CRs (left). The examples in **Fig. 5d,e and Supplementary Fig. 2b-d** each illustrate a striking correlation of both eyelid PC and MLI activity with each other and with the time of onset of the CR. Because only a subset of MLIs receive PC feedback, the model predicts that not all MLI responses should be well-correlated with CRs (**Fig. 5b2**); indeed, **Fig. 5e** shows examples with correlation coefficients ranging from 0.2 to 0.91.

We performed an analysis of the correlation between activity and CR timing for 814 MLIs (10 rabbits) during eyelid conditioning, obtained using three different ISIs. The results revealed that MLI activity could be described by two Gaussian distributions (blue and red curves in **Fig. 5b3**). The means of each subpopulation (red: r = 0.43 ± 0.01 SEM, blue: r = -0.05 ± 0.01 SEM) were similar to those predicted by the simulations that included PC feedback (**Fig. 5b2**). The skewness (rightward shift) of the *in vivo* distribution (0.84 in **Fig. 5b3**) also more closely matched the skewness (0.65) of the simulated distribution with PC feedback (**Fig. 5b2**) than that of the simulated distribution without PC collaterals (0.43; **Fig. 5b1**). Therefore, a subset of MLIs showing relatively high correlation with CRs during eyelid conditioning could be identified as putative PC-MLIs. Indeed, this analysis provides a practical tool to identify PC- MLIs *in vivo* and to investigate their functions in cerebellar learning. Taken together, our simulations and *in vivo* recordings establish that the activity of both eyelid PCs and the PC- MLIs that receive feedback inhibition from them are highly correlated with CRs during eyelid conditioning. These changes in PC and PC-MLI activity reflect plasticity within the cerebellar cortex and provide evidence that the PC-to-PC-MLI inhibitory feedback circuit plays a role in cerebellar learning.

### Activity of putative PC-MLIs during eyelid conditioning

We operationally defined putative PC-MLIs as those whose activity was correlated with eyelid position with a correlation coefficient of 0.43 or greater. This criterion identifies MLIs with a 95% or greater probability of being within the high-correlation population (red curve) of MLIs that create the high skewness of the distribution shown in **Fig. 5b3**. While this classification does not prove that such cells are PC-MLIs, our simulation results indicate that these cells are highly unlikely to be stellate cells or basket cells that do not receive PC feedback because the activity of such cells will not be correlated with CR expression. Most of the 71 putative PC-MLIs identified in this way (8.7% of all MLIs) came from sessions where activity from nearby eyelid PCs was recorded simultaneously, demonstrating that these cells were within approximately 120 µm of eyelid PCs. Correlations between the activity of all identified MLI with eyelid position and velocity are shown in **Supplementary Fig. 2a**, with the dotted line indicating the criterion level for classifying putative PC-MLIs.

Recordings from the 71 putative PC-MLIs, including the examples shown in **Fig. 5d** and the two right cells in **Fig. 5e** reveal two clear relationships between putative PC-MLI activity and eyelid CRs: *(i*) there was little or no response in these neurons in trials where no CR occurred (bottom of the sorted raster plots), and *(ii*) there was a burst of activity in putative PC-MLI that closely tracked the time of onset of CRs (red dots) on a trial-by-trial basis. **Fig. 5f** compares the mean firing rate of all putative PC-MLIs examined to the kinematics (position and velocity) of the eyelid CRs, sorted according to the ISI used for training (ISI 250 ms - blue, 500 ms - red and 750 ms - green). Despite the loss of temporal precision imposed by averaging, the firing rate of putative PC-MLIs corresponds well to both average eyelid position and velocity at each ISI. These high correlations are likely a result of activity driven by feedback from eyelid PCs, which highly anti-correlate with both measures^32^.

To quantify the temporal relationship between connected pairs of PCs and MLIs *in vivo*, we calculated cross-correlograms of MLI activity triggered on PC simple spikes. For this analysis only pairs of PCs and PC-MLIs detected by the same tetrode were used and we considered baseline activity from each unit for 10 s prior to CS onset for every trial. We were unable to evaluate the data at time 0 because simultaneously recorded action potentials from two units merge into a signal that cannot be assigned to either unit during sorting^37^. After time 0, there was a brief, significant reduction in MLI firing following a PC spike in six PC/PC-MLI pairs that were detected by the same tetrode (**Fig. 5g**; **Supplementary Fig. 3a, b**). The pattern of cross-correlation between connected PCs and PC-MLIs was similar to that observed in slices (**Fig. 4e**). As in **Fig. 4f**, this reduction in MLI activity was abolished when the timing of PC spikes was shuffled (**Supplementary Fig. 3a, b**). Differences in response time courses between *in vivo* and brain slice data are due to the lower neuronal firing rate in slices, which yields much longer time intervals between spikes, as well as temperature differences between the two experimental systems.

We also performed a similar cross-correlation analysis on PC/PC-MLI pairs in the cerebellar simulation. There was a qualitative match between the *in vivo* recordings and simulations: a transient reduction in PC-MLI firing rate was observed following a PC action potential (**Fig. 5h**) and was abolished when PC spike activity was randomized. A weaker effect of single PC spikes *in vivo*, compared to the simulations and in slices, probably arises from the larger number of spontaneous excitatory inputs to each neuron *in vivo*. In summary, cross-correlation analysis from all three data sets (**Fig. 4e** and **Fig. 5g, h**) demonstrated an inverse relationship between PC and PC-MLI activities, which is consistent with PCs providing inhibitory feedback to PC-MLIs. This correspondence strengthens the conclusion that putative PC-MLI identified *in vivo* are equivalent to the PC-MLI identified in slices.

We performed additional cross-correlation analyses for putative PC-MLI responses to either pauses or bursts in PC spiking between conditioning trials. This allowed analysis of neuronal interactions between trials, without the influence of the large anti-correlated changes in activity that occur during conditioned responses (e.g. **Fig. 5c, d**). Pauses were defined as the largest 35% of inter-spike intervals, while bursts were the smallest 35% of inter-spike intervals (50% for simulations). All spikes for a PC/PC-MLI pair were aligned to the start of the pause or burst. Pauses in PC activity clearly increased activity in putative PC-MLIs (**Fig. 5i**; average of 11/27 PC-PC-MLI pairs detected on same tetrode), while bursts of PC activity decreased putative PC-MLI activity (**Fig. 5j**; average of 13/27 PC-PC-MLI pairs). Individual examples of responses of pairs of PC and PC-MLI during PC pauses and bursts also showed strongly anti-correlated activity (**Supplementary Fig. 3c, d**). These results indicate that the feedback relationship between PCs and MLIs *in vivo* is more prominent during large changes in PC activity, rather than in response to single PC simple spikes.

To further quantify the relationship between the activity of putative PC-MLIs and the properties of the eyelid CRs, responses evoked at each ISI were grouped according to onset of the eyelid movement: early-, middle- and late-onset (**Fig. 6a**). Trials with no CR comprised a small fourth group. The timing of putative PC-MLI activity within all groups showed clear differences that closely paralleled the differences in CR latency. When trials were instead grouped according to CR peak amplitude, there were again differences in the timing of putative PC-MLI activity that correlated with differences in eyelid CR amplitude, particularly at longer ISIs (**Fig. 6b**). In summary, the activity of putative eyelid PC-MLI, like the activity of eyelid PCs^32^, is closely linked to the timing and amplitude of eyelid CRs.

**Figure 6.**
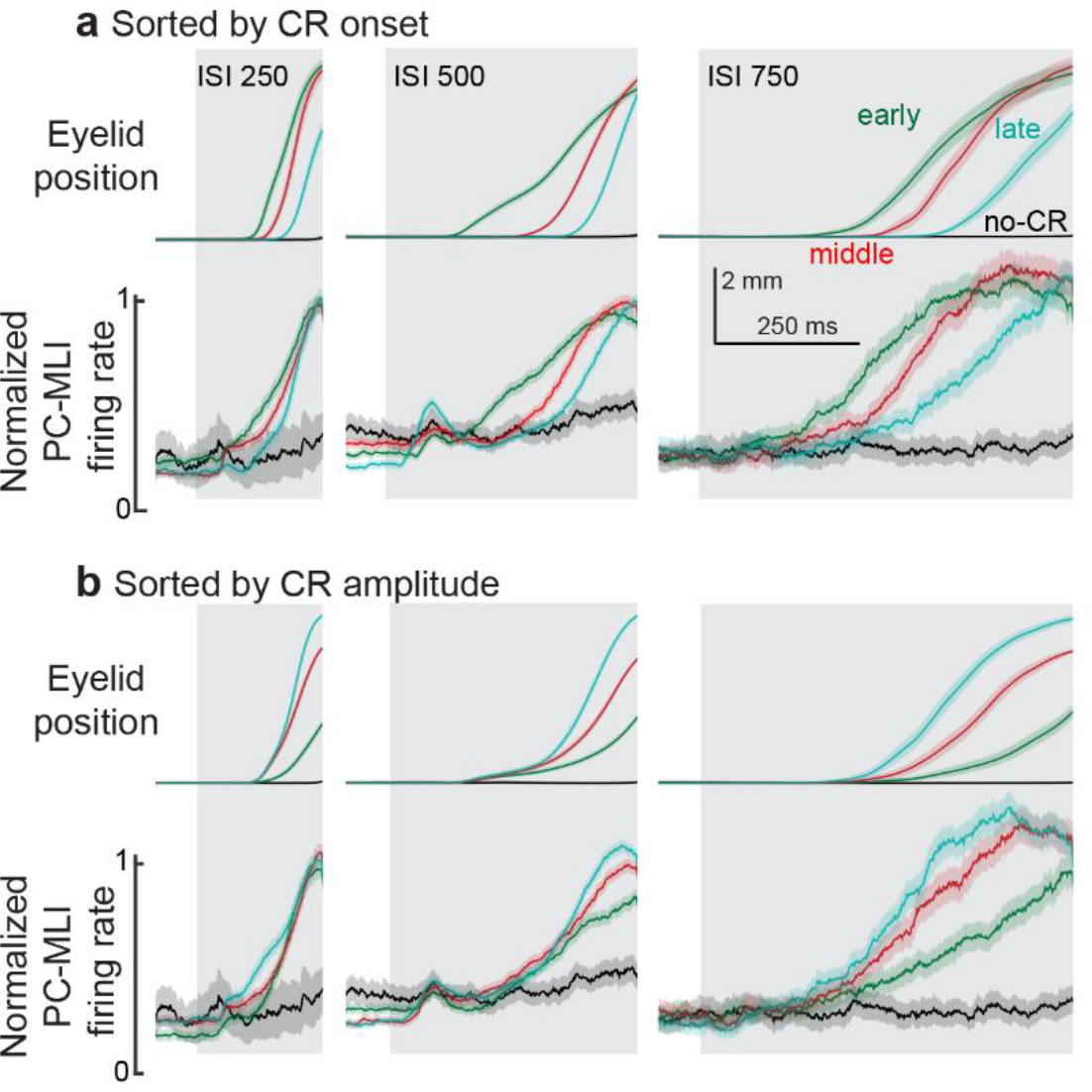
Normalized eyelid PC-MLI activity sorted by response onset or amplitude. (a) Average eyelid CRs and average responses of eyelid PC-MLIs grouped by latency to onset of the eyelid CRs. Eyelid PC-MLI activity from the corresponding groups was normalized to the maximum firing rate and averaged to determine the relationship between eyelid PC-MLI activity and differently timed eyelid CRs within each ISI. Green represents the earliest onset group, red the middle onset, cyan the latest onset and black the non-responses for each ISI. (b) Same as in **a**, e MLI t groups were sorted according to the amplitude of the eyelid CRs at the end of the ISI. Cyan represents the largest amplitude responses, red the middle amplitude, green the smallest and black the non-responses. Shaded regions represent the 95% confidence intervals.

### Temporal relationship between activity in putative PC-MLIs and Purkinje cells

Because both eyelid PC^32^ and PC-MLI responses are highly correlated with eyelid CRs (**Fig. 5c, d**), the activity of PCs and PC-MLIs also should be highly correlated during eyelid CRs. We hypothesize that the high correlation of PC-MLI activities with eyelid PCs and CRs is due to feedback inhibition from PCs to PC-MLIs, rather than feedforward inhibition from MLIs to PCs. Therefore, during eyelid conditioning, connected pairs of eyelid PCs and PC-MLIs would be expected to show anti-correlated activity, with changes in PC activity preceding those in PC-MLIs.

We tested these predictions in a subset of experiments where the activities of both putative PC-MLIs and eyelid PCs were recorded simultaneously during eyelid conditioning. Examples of such recordings, obtained at three different ISIs, are shown in **Fig. 7**. Behavioral responses (**Fig. 7a**) and recordings of neuronal activity (**Fig. 7b-d**) again were sorted according to eyelid CR latency. Peristimulus time histograms for the paired activity of an eyelid PC (green) and a putative PC-MLI (black) are shown in **Fig. 7b, c**. In each case the eyelid PC and the putative PC-MLI fired inversely, with the decrease in PC activity and the increase in PC-MLI activity both corresponding - on a trial-by-trial basis - with the eyelid CR. This inverse relationship between activity of the two cell types is most evident in superimposed plots of instantaneous firing rate for each neuron (**Fig. 7d**).

**Figure 7.**
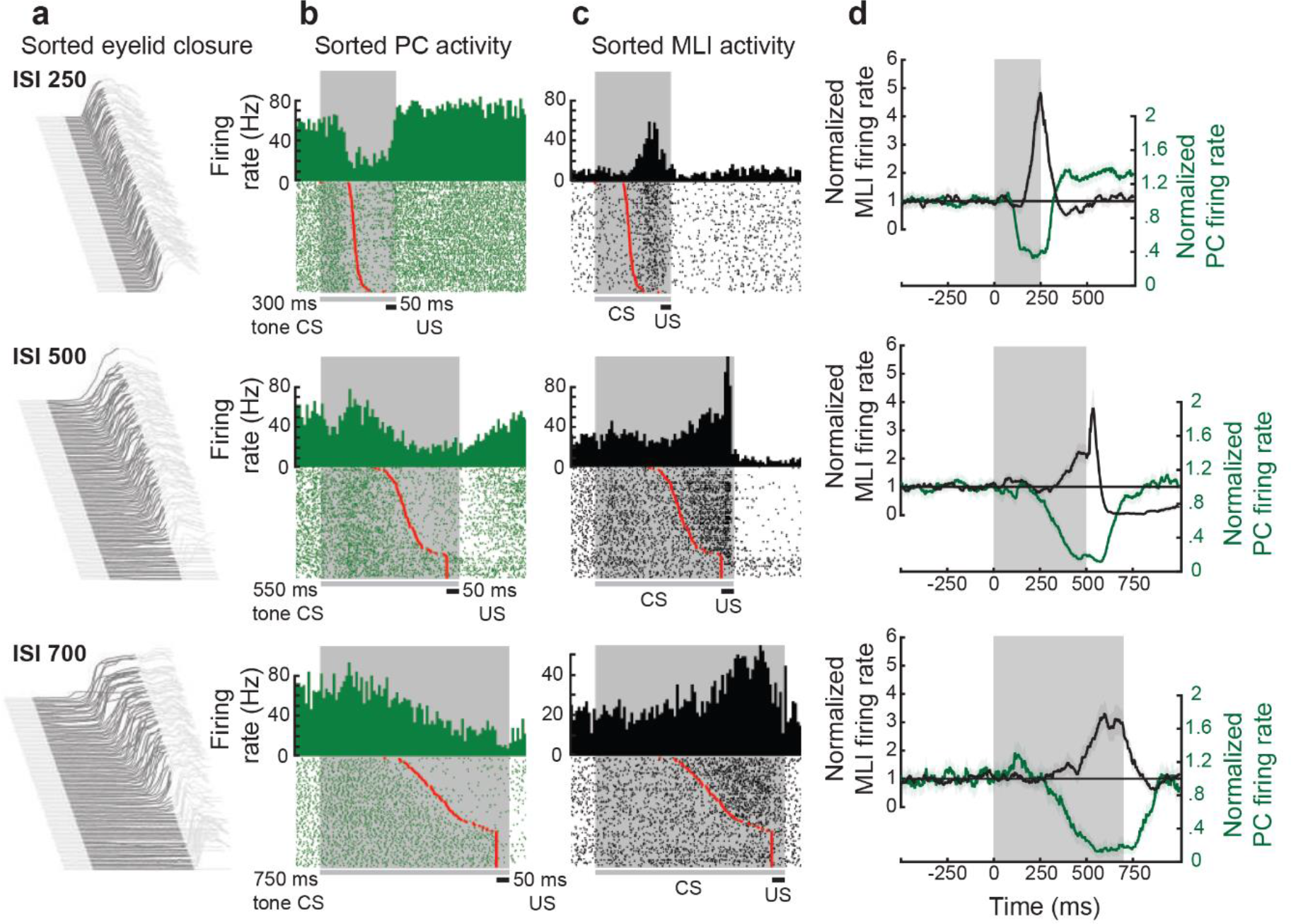
Relationship between individual simultaneously recorded eyelid MLIs and eyelid PCs during expression of conditioned eyelid responses. (**a**) Eyelid movements during an eyelid conditioning session at different ISIs, sorted according to response onset. The dark gray area indicates the duration of the ISI and upward deflection is the eyelid CR. (**b** and **c**) Peri-stimulus time histograms (10 ms bins) and raster plots sorted by response latency (red dots) of the activity of simultaneously recorded eyelid PCs and MLIs from the session in column **a**. The dark gray bars and shaded areas indicate the duration of the tone stimulus (CS) and the black bars indicate the duration of the eyelid stimulation (US). (**d**) Instantaneous firing rate for the examples in **b** and **c.** For all three examples, the activity of the eyelid PC decreases before the simultaneously recorded activity of the eyelid PC-MLI increases, as well as prior to the time of onset of the conditioned response. The gray shaded region indicates the ISI for each pair.

To examine the role of PC inhibitory feedback to PC-MLI in this correlation, we compared the predictions of computer simulations that differed only by the presence or absence of this connection. Simulations that included PC feedback inhibition of PC-MLIs predicted a strong negative correlation between eyelid PCs and putative PC-MLIs for all training ISIs (**Supplementary Fig. 4a**), while simulations lacking PC feedback showed high negative correlations at the shortest ISI, but less negative correlations at longer ISIs (**Supplementary Fig. 4b**). The same analysis of the activity of eyelid PC and PC-MLI pairs recorded *in vivo* (**Supplementary Fig. 4c**) showed a distribution quite similar to the predictions of the simulations that included PC feedback (**Supplementary Fig. 4a**). Cumulative probability plots of these data (**Fig. 8a**) revealed considerable overlap between the *in vivo* paired recordings (red) and the results of simulations including PC feedback (black), while quite different results were observed in simulations lacking PC feedback (blue). There were significant (p<0.01) differences between the *in vivo* data and the predictions of simulations without PC feedback (in 92% of bootstrap samples using two-tailed student-t test, in 86% of samples using two-tailed Kolmogorov-Smirnov test), while there were no significant differences between the *in vivo* data and the results of simulations that included PC feedback (p<0.01 in 0.1% of bootstrap samples using two-tailed student t-test, in 3% of samples using two-tailed Kolmogorov-Smirnov test; **Supplementary Fig. 4d, e**). These comparisons indicate that the anti-correlated activity of eyelid PCs and putative PC-MLI arises from inhibitory feedback from PCs to PC-MLI.

**Figure 8.**
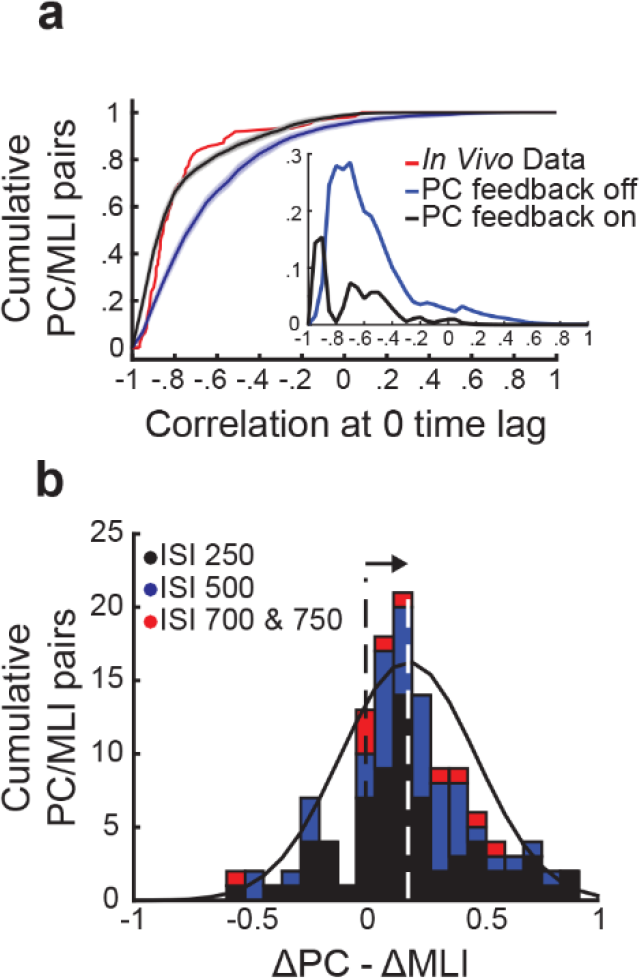
Relationship between the responses of simultaneously recorded eyelid PC- MLIs and eyelid PCs and conditioned eyelid responses. (a) Quantifying the relationship between computer simulations differing in PC feedback collaterals to MLIs with the *in vivo* paired recordings. Cumulative sum of cross-correlations from the two computer simulations and *in vivo* data along with an inset showing differences between the computer simulations both with (black) and without (blue) the feedback from PCs to MLIs and the *in vivo* recordings (red). Note how closely the *in vivo* recordings and simulation with PC feedback align, while the simulations without feedback do not. (b) Distribution showing relationship between timing of decreases in eyelid PCs activity and increases in PC-MLIs activity at each ISI. The mean of the distribution of ΔPC - ΔMLI differences is indicated with a white dotted line. This distribution is shifted to the right of 0 (arrow), indicating that the PC activity decreases more rapidly than the PC-MLI activity increases, relative to CR onset, for most simultaneously recorded pairs.

To test the prediction that inhibitory feedback from PCs to PC-MLIs causes PC activity changes to drive PC-MLI activity changes, we next quantified the temporal relationship between these two responses. At the single-trial level we measured the 50% maximum delay time (50% max response time for PC-MLI – 50% max response time for PC) between decreases in PC activity and increases in simultaneously recorded PC-MLI activity during eyelid conditioning for the pairs shown in Fig. 7 (**Supplementary Fig. 5b**).For trials with CRs, PCs reached 50% of their maximum response before PC-MLIs did (ISI 250 = 71.4 ms, ISI 500 = 80.1 ms, ISI 700 = 61.0 ms). This result is consistent with the pause/burst triggered analysis of baseline activity showing that large changes in PC activity drive opposing changes in the activity of PC-MLIs. For non-CR trials, the opposite was observed: PC-MLIs reached 50% of their maximum response before PCs did (ISI 500 = -79.3 ms, ISI 700 -70.4 ms). This reversal of timing is likely due to two factors. First, a 50% change in PC-MLIs activity requires a smaller absolute change than for PCs, because of the lower basal firing rate of PC-MLIs. The second factor is that large, learning-related decreases in PC activity were absent in non-CR trials, so the activity of both neuron types can more faithfully reflect parallel fiber excitatory inputs to both neurons.

The prediction of our hypothesis was further tested, for the entire population of PC- MLIs with simultaneously recordings of eyelid PC activity. This was done by determining a ratio representing the magnitude of changes in activity prior to CR onset, normalized to the peak amplitude of the change during the entire interval (**Supplementary Fig. 5d**). A positive difference (ΔPC – ΔMLI) indicates that the PC from each paired recording showed a larger change in activity before CR onset, while a negative difference indicates the opposite. The distribution of these differences, color-coded by ISI, is shown in **Fig. 8b**. This distribution has a mean (white dashed line) that is greater than zero (black dashed line), indicating that eyelid

PCs decreased their activity prior to an increase in putative PC-MLI activity in a majority of cases. A t-test (verified with Jarque-Bera, p = 0.5) indicated that this combined distribution was significantly different (p = 8 x 10^-15^) from a normal distribution with a mean of 0. Individual ISI distributions were also skewed rightward (skewness = .27 for ISI 250, .36 for ISI 500, and .78 for ISI 750), indicating that, on average, eyelid PC activity decreased before putative PC-MLI activity increased. Although the activity of most eyelid PCs decreased well before CR onset while PC-MLIs activity increased at or immediately after CR onset (see **Fig. 7b, c, Supplementary Fig. 5a, c, e**), in a few cases PC-MLI activity increased before eyelid PC activity decreased (**Supplementary Fig. 5e**). In these cases, the recordings may have come from neuron pairs where the PC-MLI was innervated by a different presynaptic PC.

To examine the origins of this temporal relationship between changes in eyelid PC activity (which would affect feedback inhibition) and PC-MLI activity (causing feed-forward inhibition), we tested simulations with different loci of learning-induced plasticity during conditioning. Comparisons between the predicted activity of PC-MLIs and PCs (**Supplementary Fig. 6**) indicated that the presence of plasticity at PF synapses onto PCs yielded changes in PC activity that preceded changes in PC-MLI activity, as observed *in vivo* (**Supplementary Fig. 6a, b**). However, in simulations with plasticity only at PF synapses onto MLIs, changes in firing rates of MLIs and PCs were approximately simultaneous, with changes in MLI activity slightly preceding those in PCs (**Supplementary Fig. 6c, d**). Including plasticity at both synapses produced too many MLIs that led PCs, in clear contradiction to the *in vivo* data. In summary, only simulations with plasticity exclusively at PF excitatory synaptic inputs to PCs were capable of accurately predicting our observed results.

Together, these results indicate that the observed relationships between the activity of putative PC-MLIs, eyelid PCs and eyelid CRs are best explained by the presence of a feedback circuit from PCs to PC-MLIs. This circuit forces PC-MLIs and PCs to fire inversely - with responses of most PCs leading those of most PC-MLIs - so that both eyelid PCs and putative PC-MLIs predict the features of eyelid CRs sufficiently well to be observed on a trial-by-trial basis.

### Effects of PC feedback on learning performance and circuit plasticity

To consider the behavioral consequences of the feedback circuit from PCs to PC-MLIs, we performed additional simulations, both in the presence and absence of PC feedback, that predicted the ability of the cerebellum to learn during eyelid conditioning. In these simulations, eyelid conditioning was mimicked by presenting mossy fiber and climbing fiber inputs with inter-stimulus intervals ranging from 150 to 1000 ms. Examples of virtual eyelid responses produced by simulations trained at an ISI of 200 ms are shown in **Fig. 9a**. While both simulations learned, simulations with PC feedback performed better (**Fig. 9b, c**). For example, only simulations with PC feedback were able to acquire responses at the shortest (150 and 200 ms) and longest ISIs (ISI > 700 ms), albeit exhibiting more modest learning (**Fig. 9b, c**). Even at intermediate ISIs, where simulations were able to learn without PC feedback, the presence of PC feedback improved the rate of acquisition and final asymptotic performance. This is evident from comparison of CR amplitude acquisition curves at ISI 500 ms in simulations with (black) and without (red) PC feedback (**Fig. 9b**). These results indicate that PC feedback can rapidly increase response amplitude during learning and improve performance on the temporal margins, where patterns of inputs otherwise make it difficult for the cerebellum to learn.

**Figure 9.**
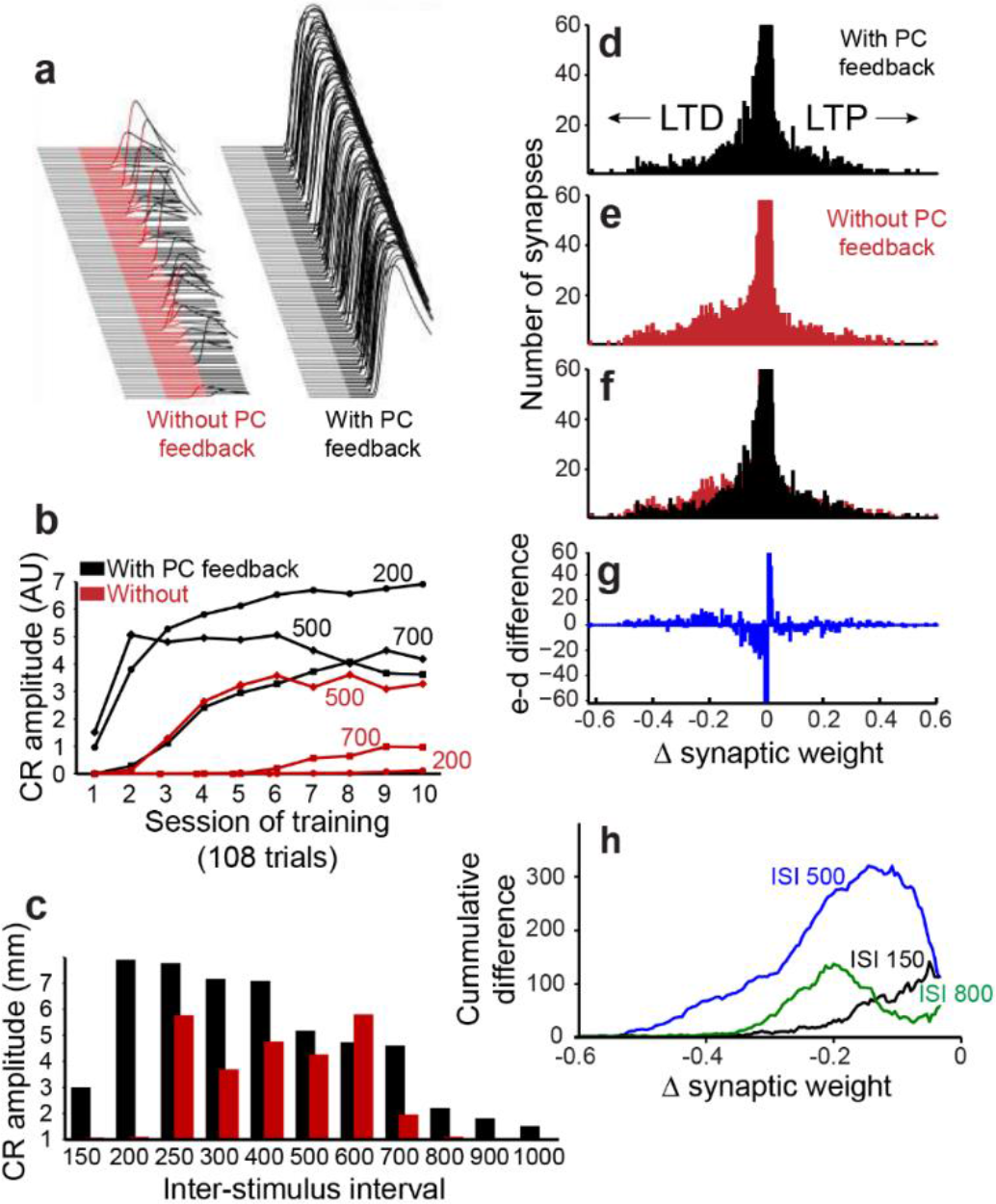
Simulation performance with and without PC collaterals. (a) Examples of virtual conditioned responses from the last session of training at ISI 200 for simulations without feedback from PCs to MLIs (CS is shown in red) and with feedback collaterals (CS is shown in black). (b) Acquisition curves for simulations constructed with (black) and without (red) feedback from PCs to PC-MLIs, plotting the amplitude of virtual CRs as a function of sessions of training for the three ISIs indicated. (c) Average virtual CR amplitude on the final session of training is shown for training using 11 ISIs ranging from 150 ms to 1000 ms; black bars indicate results from simulations with PC to PC-MLI feedback and red bars depict data from simulations lacking feedback. Simulations with PC feedback performed considerably better at relatively long and short ISIs, while for intermediate ISIs there was less effect on performance. (d) Net changes in the strength of the 12,000 synapses between granule cells and PCs in a simulation with PC feedback to PC-MLIs and trained at an ISI of 500 ms. Negative changes indicate net LTD (left), while positive changes indicate net LTP (right). The vertical scale of the histogram is cropped at 60 synapses because the vast majority of synapses showed little or no change. The synapses that underwent net LTD are most likely to support expression of the conditioned responses elicited by the simulations. (e) Same analysis as in **d**, for a simulation lacking PC feedback to PC-MLIs. (f) Superimposition of **d** and **e** indicates a larger number of synapses underwent robust LTD in the absence of PC feedback. (g) Difference between **d** (with feedback) and **e** (without feedback). The simulation lacking PC feedback to PC-MLIs had a larger number of synapses undergoing robust LTD and fewer synapses undergoing a small degree of net LTD. (h) To compare the differences between simulations with and without PC feedback, across several ISIs, difference plots such as that shown in **g** were converted to a cumulative histogram. Data from ISI 150, 500 and 800 ms are shown. Across these ISIs, the simulations lacking PC feedback collaterals had more synapses undergoing robust LTD than those with feedback. This was true for ISIs where the simulations learned approximately equally well (ISI 500 in blue) and for ISIs where simulations with collaterals learned much better (ISIs 150 and 800).

To better understand how PC feedback improves learning, we used the simulation results to calculate training-induced changes at all 12,000 PF excitatory synapses between granule cells and PCs, which are predicted to be primarily responsible for learning. In **Fig. 9d- f**, synapses that underwent net LTD appear as negative values, while those showing LTP have positive values. While most synapses underwent little or no net change in strength, in both simulations more PF synapses showed LTD than LTP (**Fig. 9d, e**). However, the presence of PC feedback was predicted to yield robust decreases in strength at fewer synapses (net LTD; **Fig. 9f**); subtraction of the results from the two simulations reveals the net decrease in learning-dependent LTD of PF synapses produced by PC feedback (**Fig. 9g**).

To compare this effect across all 11 ISIs, we converted difference histograms, such as the one in **Fig. 9g**, into cumulative distributions of changes in synaptic weights (**Fig. 9h**); the blue line (ISI 500) is derived from the data shown in **Fig. 9g**. For all ISI, PC feedback is predicted to cause fewer synapses to undergo LTD (**Fig. 9h**). In summary, simulations with PC feedback learned better and required less net LTD of synaptic efficacy, indicating more efficient distribution of learning-induced changes in synaptic strength.

## DISCUSSION

We used optogenetic circuit mapping and paired electrophysiological recordings in cerebellar slices to demonstrate functional inhibitory connections between PCs and PC-MLIs and to characterize the physiological properties and spatial organization of this novel synaptic circuit. Our optogenetic circuit mapping enabled the first detailed analysis of the function and spatial organization of this circuit. A large-scale simulation of the cerebellum suggested that this interconnectivity between PCs and PC-MLIs increases cerebellar efficiency by permitting learning with less overall net synaptic plasticity. Finally, we used *in vivo* recordings from MLIs during delay eyelid conditioning, a well-established cerebellum-mediated task, to demonstrate predictions of the simulation that the activity of some MLIs was correlated with the behavioral conditioned responses and anti-correlated with the activity of eyelid PCs.

### Functional organization of Purkinje cell-MLI circuits

It has long been known that PC collateral axons bifurcate and extend along the PC layer^5, 38–41^, to inhibit other PCs and to reach into the lower molecular layer and upper granule cell layer. In rodents, PC axon targets outside of the PC layer are mainly MLIs within the molecular layer as well as Lugaro cells and granule cells in specialized regions of the granule cell layer^5, 38, 40, 42^. Here we build upon this anatomical information, and limited physiological analysis done *in vivo*^4, 43^ or in slices^5^, to functionally characterize the monosynaptic connection between PCs and MLIs.

Connections between PCs and MLIs are selective: PCs apparently connect to only a fraction of MLIs within the inner one-third of the molecular layer. It remains to be determined whether the MLIs that receive PC feedback inhibition are basket cells or some other type of interneuron. Our connectivity rate is lower than a previous observation^5^; this difference may arise from our more reliable (seven times larger) sample size. Although the fraction of MLIs receiving PC input could be underestimated, due to loss of connections during slicing of the cerebellum, our observations are consistent with anatomical observations of sparse representation of PC axons within the molecular layer^9, 38^. Our observation of no functional connectivity between PCs and stellate cells in the outer two-thirds of the molecular layer is also consistent with anatomical studies^39, 44^. These connectivity features account for the difficulty in finding connected pairs of neurons, both *in vitro* and *in vivo*.

We were able to characterize the fundamental properties of the PC-to-PC-MLI circuit. Inhibitory synaptic responses evoked by PC stimulation were sufficiently strong to prevent action potential firing in postsynaptic PC-MLIs. These responses had a short onset latency and were unaffected by a glutamate receptor antagonist, consistent with a monosynaptic connection. Other noteworthy properties of IPSCs at this synapse include variable amplitudes and fast rise times. These properties generally are similar to those of IPSCs at the inhibitory synapse formed between PCs via axon collaterals^5, 9, 12^, aside from a lower rate of failure at the PC-to-PC-MLI synapse. The latter property arises from a relatively large readily releasable pool of synaptic vesicles, as well as a relatively low release probability. A remarkable activity-dependent enhancement of mobilization of reserve pool synaptic vesicles allows this synapse to sustain transmission even at the high rates of activity typical of PCs.

Our high-speed mapping visualized the spatial organization of PC inputs onto a postsynaptic MLI and revealed one or two discrete areas of input within the molecular and PC layers, where PCs are located. The presence of PC-MLI receiving two discrete areas of presynaptic PC input (e.g. **Fig. 3d2**), as well as a mean presynaptic input area 1.5 times larger than the size of the optical footprint of a single PC (**Fig. 3e**), indicate that on average one or two PCs converge onto a postsynaptic MLI. This is in line with the relative abundance of PCs and PC-MLIs: our measurements indicate that approximately 1/18 of all MLIs receive PC feedback, while previous estimates suggest that the number of PCs is approximately 1/10 that of MLIs. Thus, PCs are 1.8 times more abundant than PC-MLIs, which is in reasonable agreement with the 1.5-fold convergence between PCs and PC-MLI that we observed. Because PCs are the sole output of the cerebellar cortex, any cell that influences PCs will influence behavior; the similar numbers of PCs and PC-MLIs indicates high potential for such influences. The relative abundance of PCs and PC-MLI also suggests that nearly every PC provides feedback inhibitory drive to PC-MLIs. Our observation that PC-MLIs sometimes receive two convergent PC inputs contrasts with a previous suggestion that only single PCs innervate MLIs^4^. PC-MLI connections were mostly located within 200 µm from postsynaptic PC-MLI somata, consistent with the reported sagittal extent of PC axon collaterals^5, 9, 41^. We also found that the majority of presynaptic PCs were located on either the apical or juxtaposed side relative to postsynaptic MLIs, verifying anatomical observations^5, 9^.

Despite using two experimental strategies that optimized detection of either PC-to-MLI or MLI-to-PC connectivity, in no case were reciprocally connected pairs of PCs and MLIs observed (**Fig. 4c**). Although it was previously assumed that PC-to-PC-MLI circuits are non-reciprocal^4, 42^, our results provide the initial experimental test of this important point. Such an arrangement allows feedback inhibition of PC-MLIs by PCs to disinhibit neighboring PCs (**Fig. 4d**), which may help to synchronize PC activity in the sagittal plane. It also will serve to prevent network oscillations, which are often the outcome of reciprocal inhibitory circuits^45^.

### *In vivo* activity of MLIs

This interconnectivity between PCs and MLIs gave rise to specific and testable predictions about the activity of PC-MLIs *in vivo*. MLIs that receive input from PCs should fire inversely with PCs (**Fig. 4e**) and, like PCs, their activity should correlate on a trial-by-trial basis with cerebellar-mediated behavioral responses. We tested these predictions using eyelid conditioning because: (1) the role of the cerebellum in this task is well established; (2) the activity of eyelid PCs is well characterized and appears to control the timing and amplitude of the behavioral conditioned responses; and (3) computer simulations of the cerebellum allowed us to explore the plausibility of alternative explanations for why MLI activity correlates, on a trial-by-trial basis, with PC activity and behavioral responses.

Recordings from MLIs during expression of conditioned eyelid responses supported these predictions. Some MLIs showed unusually high trial-by-trial correlations with the eyelid responses. Computer simulations predicted such high correlations only when PC-to-PC-MLI connections were included, even when these simulations were constructed to exaggerate contributions from other sources. Simultaneous recordings showed that changes in eyelid PC activity tended to precede, on a trial-by-trial basis, that of putative PC-MLIs (**Figs. 7b-d** and **8b**). Cross-correlation analysis revealed similar inverse activity correlations between PCs and PC-MLIs across all three platforms we examined: cerebellar slices (**Fig. 4e**), *in vivo* recordings (**Fig. 5g**) and computational modeling (**Fig. 5h**). Finally, we observed that computer simulations trained with eyelid conditioning procedures learned better when PC-to-MLI connections were included, despite these connections yielding less net synaptic plasticity (**Fig. 9**). Even though previous versions of our simulation that did not include PC-to-MLI feedback could exhibit learning^20, 21^, simulations that included PC-MLI connections performed better, particularly in difficult learning conditions.

These results suggest that in networks where PCs inhibit MLIs, each PC acts to make other PCs fire more like itself through disinhibition (**Fig. 4d**). In this way, PC-MLIs can exert strong influence over cerebellar processing and learning despite being the least abundant MLI. Because PC-MLIs are both inhibited by PCs and inhibit other PCs, their connectivity confers upon them particularly strong influence over PC activity and thence over the output of the cerebellar cortex. We observed inverse activity between PCs and PC-MLIs during instances of large increases or decreases in PC activity, both during large pauses in PC activity controlling expression of conditioned responses and during the largest changes in PC activity during the inter-trial interval (pause/burst triggered averages). As a consequence, the function of PC feedback to PC-MLIs could be to selectively communicate large changes in PC activity within a parasagittal strip through these PC-MLIs. Inhibitory neurons are ubiquitous throughout the brain and are often connected to each other by inhibitory synapses. Therefore, our results suggest a computational principle: activation that involves disinhibition – as occurs between PCs and certain MLIs – could synchronize the activity of inhibitory neurons. Here, we use “synchrony” as a broad term that includes PC pause synchronization. Pause synchrony can be exerted as diverse forms (pause beginning, pause ending, pause overlapping synchrony) for several milliseconds, which results in a variety of forms of modulation of neuronal activity and may lead to a broad range of behavioural outcomes^46, 47^.

One apparent consequence of disinhibition-based synchronization of learning-related activity in PCs suggested by our simulations is to require a smaller net change in excitatory synaptic input for PCs to decrease their activity to the level required to produce a well-timed CR during the CS. Such economy of plasticity could be greatly beneficial for faster learning, better learning at difficult ISIs, and increased capacity to store learned responses. Strictly speaking, plasticity at PC synapses cannot be inferred from learning-related changes in PC activity: these changes could instead result from plasticity at synapses onto MLIs. While it is well-established that plasticity occurs at PF synapses between GCs and PCs^33^ (blue in **Fig. 5a**), plasticity at GC-MLI synapses (green in **Fig. 5a**) has only been inferred from learning-related changes in MLI activity and from MLI receptive field expansion^48, 49^. Previous work showed evidence of learning-dependent changes in MLI activity that corresponded to CRs during eyelid conditioning in mice, which was also interpreted to result from plasticity at MLI synapses^50^. Our results substantially limit such inferences by showing that learning-dependent changes in MLI activity could instead come from interactions between PCs and PC- MLIs. While our results do not completely exclude the possibility of learning-related changes at PF synapses innervating MLIs, they do demonstrate that learning-related changes in MLI activity (or PC activity) do not necessarily indicate changes in the strength of synapses onto MLIs (or PCs). Further studies will be required to fully sort out the possible contributions of MLI synaptic plasticity to learning. Independent of detailed mechanism, we nonetheless conclude that feedback inhibition from PCs to PC-MLIs contributes to more efficient cerebellar-mediated motor learning.

## METHODS

### In vitro electrophysiological recordings and optogenetic mapping

*Mice.* PCP2 (Purkinje cell protein 2 or L7)-ChR2-H134R mice were generated as previously reported. PCP2-cre transgenic mice [(Pcp2-cre)2Mpin/J or (Pcp2-cre)3555Jdhu/J; Jackson Labs]^6, 8^ were crossed with mice expressing floxed ChR2-H134R [B6;129S- Gt(ROSA)26Sor^tm32(CAG-COP4*H134R/EYFP)Hze/J^]^7^. Mice positive for both transgenes were selected through PCR-based genotyping^16^ using the following primers: forward primer; 5’-GCG GTC TGG CAG TAA AAA CTA TC-3’ and reverse primer; 5’-GTG AAA CAG CAT TGC TGT CAC TT-3’^8^. Mice were maintained under a 12h light/dark cycle with free access to food and water. All procedures were conducted according to the Institutional Animal Care and Use Committee guidelines of the Biopolis Biological Resource Center.

### Brain slice recording

Conventional methods were used to prepare cerebellar slices from mice aged between 14 days to 77 days^16, 17^. Mice were sacrificed by decapitation under deep halothane anesthesia and the brain was rapidly removed. Sagittal cerebellar slices (300 µm thickness) were cut with a Leica vibratome in ice-cold oxygenated cutting solution containing (in mM): 250 sucrose, 26 NaHCO3, 10 glucose, 4 MgCl2, 3 myo-inositol, 2.5 KCl, 2 sodium pyruvate, 1.25 NaHPO4, 0.5 ascorbic acid, 0.1 CaCl2, 1 kynurenic acid. After cutting, slices were kept for at least 1 hr in the oxygenated standard external solution which contained (in mM): 126 NaCl, 24 NaHCO3, 1 NaH2PO4, 2.5 KCl, 2.5 CaCl2, 2 MgCl2, 10 glucose, 0.4 ascorbic acid (pH 7.4 when equilibrated with 95% O2/5% CO2). In some experiments, bicuculline (10 μM; Sigma), GABAzine (SR-95531, 5 μM; Sigma) or kynurenic acid (2 mM; Sigma) was added to the external solution to block chemical synapses.

Whole-cell patch clamp recordings were performed at room temperature while the slices were in the recording chamber continuously perfused with extracellular solution. Patch pipettes for Purkinje cells (3-6 MΩ) and MLIs (6-10 MΩ) recordings were pulled on a PC-10 puller (Narishige, Japan). Pipettes were filled with a solution containing (in mM): 130 K-gluconate, 10 KOH, 2.5 MgCl2, 10 HEPES, 5 EGTA, 4 Na2ATP, 0.4 Na3GTP, 5 disodium phosphocreatine (pH 7.3, ∼295 mOsm). Membrane potentials were not corrected for liquid junction potentials. The IPSC reversal potential was -74 mV (n = 3). Alexa 594 (50 μM) was added to the internal solution to image cell morphology. In a subset of interneuron recordings, QX-314 (10 mM; Sigma) was included in the internal solution to block MLI action potentials. IPSCs were recorded under voltage clamp at a holding potential of -40 mV, unless otherwise indicated. Electrical responses were acquired via a Multiclamp 700B amplifier (Molecular Devices) and digitized at 100 kHz via a Digidata 1440A interface (Molecular Devices).

To measure synaptic depression, three different stimulation methods were used: dual electrophysiological recordings, optogenetic stimulation and extracellular stimulation. Because photostimulation could only evoke PC activity up to 25 Hz^16^ and paired PC-PC-MLI recordings were difficult to obtain, we primarily performed extracellular stimulation. To do this, a glass pipette filled with external solution was placed in the granule cell layer below the presynaptic PC area identified by optogenetic mapping. Experiments were done at 34–35°C, to more closely emulate physiological conditions, and kynurenic acid (2 mM) was used to block glutamatergic input to PC-MLIs.

### Photostimulation

Photostimulating Purkinje cells was accomplished as previously described^17^. For wide-field excitation, blue light (465-495 nm) illuminating relatively large area (∼0.233 mm^2^) on brain slices was provided by a mercury arc lamp and light pulse duration was controlled by an electronic shutter (Uniblitz T132; Vincent, Rochester, NY). To perform high-speed circuit mapping, a laser-scanning microscope (FV1000MPE; Olympus, Tokyo, Japan) equipped with × 25 NA 1.05 (Olympus XLPlan N) water-immersion objective lens was used to create small spots of laser light to photostimulate ChR2-expressing Purkinje cells. A 510 x 510 µm area of the slice was scanned with a 405-nm laser spot (4 ms duration) in a 32 x 32 array of pixels, yielding a scanning resolution of 16 µm. The laser spot was scanned in a pseudorandom sequence, to avoid photostimulation of adjacent pixels, while cellular responses were simultaneously measured in whole-cell patch clamp recordings. The same microscope was used for confocal imaging of neuronal structure with Alexa 594.

In the course of our experiments, we used approximately 140 PCP2-Cre Mpin mice. In 4 of these mice (3%), we saw expression of ChR2 in MLI (and in other cells beyond Purkinje cells), readily evident as measurable photocurrents in response to light stimulation, consistent with a previous observation^5^. These mice were discarded and were not included in our analyses. Even in these 4 mice, the laser intensity required to evoke action potentials in MLIs non-specifically expressing ChR2 was greater than 6 µW; because we used a laser intensity of 3 µW in our mapping experiments, there was no chance of evoking action potentials in MLI in any case. Amongst the remaining 136 mice that we did use for our experiments, we recorded from more than 400 MLI and never detected measurable photocurrents. Thus there was no ChR2 expression in MLI under our experimental conditions. Furthermore, the lower connectivity rates in our results, compared to that measured previously^5^, and the restriction of synaptic responses to MLIs within inner molecular layer, not stellate cells, both indicate that ectopic expression of ChR2 in MLIs is not an issue in our experiments. The size and location of the input maps evoked by light in our mapping experiments more closely resemble the optical footprints of PCs (Fig. 2a) than MLI^17^ further suggests that all light-evoked IPSCs that we measured were a result of PC inputs rather than MLI inputs, which have not yet been found between PC-MLI and other MLIs. We also obtained similar results in another PCP2 -cre line^5^, further indicating that our optical circuit mapping precisely detected the connectivity between PCs and PC-MLIs.

### Data analysis

Mapping data were analyzed with custom software written in MATLAB by P. Namburi as previously described^17^. Spatial maps of the light sensitivity of ChR2-expressing Purkinje cells (optical footprints) were created by correlating the location of the photostimulation spot with action potentials evoked between 0-14 ms after the start of a light pulse (4 ms duration). Due to the relatively high spontaneous activity of Purkinje cells, typically 2 or 3 maps were averaged to reduce the influence of this background activity: only locations where action potentials were consistently evoked were included within optical footprints.

To generate synaptic input maps, the location of the light spot was correlated with the resulting IPSCs in molecular layer interneurons. The minimum threshold amplitude for detecting IPSCs was 15 pA. The relatively high rate of spontaneous IPSCs created background noise; this was reduced by averaging together 2-3 input maps, with pixels having an IPSC detection probability of 0.5 or greater considered to arise from photostimulation of presynaptic PCs, rather than spontaneous activity. The area of the input field was measured by summing all pixels showing suprathreshold responses. Single-pixel areas within input maps also result from spontaneous PC activity and also were excluded from analysis^17^.The peak amplitude and latency of IPSCs were determined from the maximum IPSC measured between 4 and 24 ms after the start of the light pulse. Onset times were estimated as the zero-crossing time of a line joining values measured from 20% to 80% of the IPSC peak^49^. Data are expressed as mean ± SEM.

### *In vivo* recordings and behavioral training methods

#### Rabbits

Ten male New Zealand albino rabbits (Oryctolagus cuniculus; Myrtle’s Rabbitry), weighing 2.5-3 kg at experiment onset, were used for *in vivo* experimentation. Treatment of rabbits and surgical procedures were in accordance with National Institutes of Health guidelines and an institutionally approved animal welfare protocol. All rabbits were maintained on a 12 h light/dark cycle.

#### Surgery

One week before the start of recording, rabbits were removed from their home cage and anesthetized with a cocktail of acepromazine (1.5 mg/kg) and ketamine (45 mg/kg). After onset of anesthesia, animals were placed in a stereotaxic frame and maintained on isoflurane (1∼2% mixed in oxygen) for the remainder of the surgery. Under sterile conditions the skull was exposed with a midline incision (∼5 cm), and four holes were drilled for screws that anchored the head bolt in place. The animal’s head was then positioned with lambda 1.5 mm ventral to bregma and a craniotomy was drilled out at 5.9 mm posterior and 6.0 mm lateral to lambda. The skull surface was marked and skull fragments were carefully removed from the craniotomy along with the dura matter under visual guidance. A custom-made hyperdrive array (12 or 16 tetrodes, 2 references) fitted with an electronic interface board (EIB-54 or EIB36- 16TT, Neuralynx) was implanted in the left anterior lobe of the cerebellar cortex at 17.8 mm ventral to lambda. Final ventral placement of tetrodes during surgery was between 1 to 2 mm above the target coordinate to allow advancement of the tetrodes to the target. Hyperdrives were positioned at a 40° angle caudal to vertical to avoid the cerebellar tentorium. This region of the cerebellum has been shown to be involved in acquisition and expression of well-timed conditioned eyelid responses in rabbits^28^. The bundle cannula of the hyperdrive was lowered to the surface of the brain and the craniotomy was sealed with low viscosity silicon (Kwik-Sil; World Precision Instruments). A screw attached to an insulated silver grounding wire (.003” bare, .0055” coated, A-M Systems) was then screwed into the skull. The silver wire was also attached to the ground channel of the EIB with a gold pin. The head bolt, screws, and hyperdrive were secured with dental acrylic (Fastray Pink; The Harry J. Bosworth Company), and the skin was sutured where the skull and muscle was exposed. Finally, two stainless steel stimulating electrodes were implanted subdermally caudal and rostral to the left eye. Rabbits were given analgesics and antibiotics for two days after surgery and monitored until fully recovered.

#### Tetrode recording and unit isolation

Each independently movable tetrode was composed of four nichrome wires (12 µm diameter; Kanthal Palm Coast), twisted and partially melted together to form a tetrode. Individual wires of each tetrode were connected to the EIB with gold pins, all four wires of the reference tetrodes were connected to a single reference channel of the EIB. Each tetrode was gold plated to reduce final impedance between 0.5 to 1.5 MΩ measured at 1kHz (impedance tester IMP-1; Bak Electronics). Tetrodes targeting cerebellar cortex were placed over the left anterior lobe and advanced to within 2.0 mm of the target during surgery using stereotaxic guidance. Tetrodes were then lowered in 40 to 80 µm increments during turning sessions (∼ 1 hour) until at least one stable single unit was identified; there were often multiple units on a single tetrode. After turning sessions, tetrodes were allowed to stabilize for at least 24 h and units were checked again before recording and behavioral training commenced. Putative PCs were initially identified by their higher baseline firing rate relative to cerebellar cortical interneurons and later confirmed by identifying complex spikes during cluster cutting^32^. MLI single units were often recorded simultaneously on the same or different tetrodes during recordings from PCs. Recordings were done once a single unit was identified and stable without knowledge of the origin of its climbing fiber input for PCs or identity of MLI type. Tetrodes in cerebellar cortex targeted the region containing PCs receiving evoked complex spikes from eyelid stimulation (**Supplementary Fig. 1a**). The use of tetrodes allowed for simultaneous recording of multiple single units, including MLIs, during conditioning experiments. All PCs included in the analysis had confirmed spontaneous complex spikes and all single units were held throughout the entire session. Any recording that was lost during a recording session was not included in the analysis.

Neuronal signals were first preamplified at unity gain. The signals were then fit to a window between 250 to 2000 µV and bandpass filtered (0.3-6 kHz; Neuralynx). Neural signals that exceeded a channel amplitude threshold were digitized and stored at 32 kHz (Cheetah system; Neuralynx). Custom interactive cluster cutting programs were used to manually isolate and identify single units (**Supplementary Fig. 1b**). Waveform characteristics were plotted as a two-dimensional scatter plot of one wire of the tetrode versus another in terms of energy, peak, and valley measures. The energy measure represents the square root of the sum of the squared points for the entire waveform. The peak measure is the maximum height (positive amplitude) of the waveform. The valley measure is the maximum depth (negative amplitude) of the waveform. When possible the initial identification of a single unit was made using peak as channel thresholds were set during recordings with that feature of the waveform. For recordings in cerebellar cortex a late peak measure, defined as the maximum peak during the last five points of the thirty-two points that make up each waveform, was also used to identify the later peak component of the complex spikes from the earlier peak of the simple spikes. From these cluster cutting analyses, a single PC recording would then yield two clusters, one containing simple spikes cut using peak, valley, and energy, and the second containing the complex spikes cut using the late peak parameter^32^. Following cluster cutting, all subsequent data analysis was performed using custom-written scripts in MATLAB.

#### Conditioning procedure

Conditioning experiments and recordings were done in custom training chambers (89 X 64 X 49 cm). Rabbits were placed in a plastic restrainer and the ears were stretched over a foam pad and taped down to limit head movement. An adjustable infrared emitter/detector was secured in place with the head bolt and aligned to the middle of the left eye. The infrared emitter/detector measured eyelid position by converting the amount of emitted infrared light reflected back to the detector, which increases as the eyelid closes, to a voltage. The signal was amplified to yield a signal that was linearly related to upper eyelid position (± 0.1 mm). The eyelid position detector was then calibrated before each training session by delivering a test US to elicit maximum eyelid closure (6.0 mm). The corresponding voltage deflection (∼ 6V) was then divided by 6 mm to obtain a mm/V calibration. Each training chamber was also equipped with a speaker connected to a stereo equalizer and receiver which were connected to a computer that generated the tone conditioned stimulus (CS). The CS used during training was either a 1 or 9.5 kHz sinusoidal tone (85 dB), which ramped at onset and offset with a 5 ms time constant to avoid audible clicks from the speaker. To deliver the US, leads from a stimulus isolator (Model #2100, A-M Systems) were attached to electrodes caudal and rostral to the eye. The US was eyelid stimulation, which consisted of trains of 1 ms current pulses delivered at 100 Hz for 50 ms. The intensity was adjusted for each animal to be just above threshold to elicit a full eyelid closure (between 0.8 and 1.5 mA, depending on the condition of the implanted wires).

Stimulus presentation was controlled by custom software operated on a Windows XP-based computer. To permit temporal alignment of neural and behavioral responses, digital timing pulses were generated by the computer controlling stimuli and measuring behavior and were sent to the digital input port on the Digital Lynx acquisition system (Neuralynx). During initial paired delay conditioning the tone CS was 550 ms which co-terminated with the 50 ms eyelid stimulation US which produced an ISI of 500 ms. All rabbits were initially trained and extinguished with delay conditioning at ISI 500. Further conditioning sessions involved either a 1 or 9.5 KHz tone at ISIs of 250, 700 or 750 ending with the same 50 ms eyelid stimulation US. Each training session consisted of twelve nine-trial blocks (108 trials) with each block starting with a CS alone trial followed by eight paired CS-US trials. The mean intertrial interval was 30 s with a range of 20 to 40 s.

#### Eyelid position analysis

For each trail, 2,500 ms of eyelid position (200 ms pre-CS, 2,300 ms post CS) were collected at 1kHz and at 12 bit resolution. Data were stored to a computer disk for subsequent off-line analysis. Eyelid position data was passed through a low-pass filter. Response measures calculated for each trial included CR amplitude, latency to CR criterion and latency to CR onset (**Supplementary Fig. 1c**). CR amplitude was defined as the value of eyelid position from the baseline at the time of US onset. Latency to CR criterion was defined as the time point at which the CR reached the .3 mm criterion to be designated as a CR. Latency to CR onset was determined using a custom-written two-step algorithm. The first step was designed to detect the initial deflection away from the pre-CS baseline, while the second step used linear interpolation to determine the exact time of CR onset. For further analyses eyelid trajectories were truncated at US onset to exclude non-cerebellar influence on the eyelid movement.

#### Single unit recording analysis

Instantaneous firing rate of each single-unit recording was estimated on every trial using a one-sided Gaussian kernel with a 25 ms standard deviation window. We chose a one-sided Gaussian to prevent neural responses related to the US from contaminating unit activity during the CS. PCs firing rate was normalized by the value of the baseline firing rate during 1500 ms of pre-CS activity. Each MLI firing rate was normalized, however, to the maximum during the CS of the average firing rate through the session. Most putative PC-MLI had a similar baseline firing rates, but showed variable CR-related increases in activity, ranging from two-fold increases to eight- to ten-fold increases over baseline firing rates. Under these conditions, calculating the simple mean of all PC-MLI activity or normalizing it to the baseline firing rate would yield overrepresentation of PC-MLIs with the highest CR-related increases in activity. To avoid such bias, we normalized PC-MLI responses to the maximum mean firing rate during the CS.

For each MLI single unit, cross-correlations between firing rate and behavioral responses were calculated on every trial when the animal produced a CR. For every trial, starting from 150 ms before CS onset and through US onset, we calculated cross-correlations between the instantaneous firing rate profile and the time profile of the CR (eyelid position). Then, cross-correlation values were averaged through CR trials to produce an average single-trial correlation with behavior for a given MLI (see **Fig. 5**). For comparison of activity of simultaneously recorded eyelid PCs and MLIs, cross-correlations were calculated between firing rates then averaged through the session (see **Fig. 8**). In all cases, mean values of arguments were subtracted before calculating the correlation value.

To address the temporal relationship between the activity of PCs and PC-MLIs, we developed the following analysis; this used only sessions with simultaneously recorded eyelid PCs and PC-MLIs. First, since both PC and MLI activity is precisely temporally connected with behavior, we calculated average firing rates, aligning each trial to the time of CR. This procedure reduced the influence of CS related changes in activity on the results, phasic responses to the CS for example. Depending on ISI, we used either 100 ms (ISI 250) or 150 ms (ISIs 500, 700 and 750) after CR onset time. Trials with CR onsets occurring earlier that these values before the US were excluded from the analysis, as they would have contaminated the results with US related responses. Second, for both PCs and MLIs we calculated the fraction of the full decrease or increase respectively that happened prior to CR onset. A value of 0 with this analysis indicates that there was no change in that cell’s response prior to CR onset, while a value of 1 indicates that the maximum change in firing happened prior to CR onset. Due to this measure being dimensionless it provides the same type of normalization for both cell types, thus, we used it to find whether PCs or MLIs typically change their activity first before CR onset.

For spike-triggered cross-correlogram analysis we aligned spike times of unit 1 within a +/- window onto the spike times of unit 2 (1 ms bins, ± 30 ms). The process was repeated for all spikes of unit 2 during 10 seconds before each trial onset, to produce a spike count of unit 1 as a function of time. The significance of modulation in cross-correlograms was assessed by performing the same analyses on shuffled spike times, making unit 1 and unit 2 independent, and computing the mean and standard deviation of cross-correlogram spike count of shuffled data. Normalized spike count, shown on Figure 4 and Suppl. Figure 4, was computed by dividing spike count of unit 1 by the mean of the shuffled data. Z-scores were then computed to identify significantly modulated pairs of PCs and MLIs. Significantly modulated pair was defined as having at least 2 time-bins, excluding t = 0 bin, reaching above 95% significance thresholds (Z = ± 3.34, corrected for multiple comparisons with Bonferroni correction). The procedure of the pause/burst triggered cross-correlogram analysis was similar to the spike triggered, except data was aligned only to the largest (or the smallest) 35% (50% for simulation) of PC inter-spike intervals, which were operationally defined as pauses and bursts of PCs activity respectively. Significantly modulated pairs were defined as having by at least 2 significant bins more after t = 0 compared to before t = 0 epoch.

#### Grouping

To further investigate how MLIs correspond to the onset and amplitude of conditioned eyelid responses, data from each ISI were divided into equal subgroups of trials by CR onset in relation to CS onset or by CR amplitude. Single-unit data were divided with respect to CR onset and amplitude in order to further differentiate unit activity during CRs with different onsets and amplitudes due to the differing amount of variability observed in each response measure. For each ISI, the data for each subgroup was divided equally so the same percentage of trials exists within each CR onset range or amplitude range. A few exceptions to dividing the data within each ISI equally were unavoidable and mostly involved the percentage of data in the group representing the non-CR trials within each ISI. Dividing the data with respect to CR onset involved aligning each trial by CS onset and sorting the behavioral data into equal groups with respect to the CR onset time. Subgroup eyelid CRs and MLI data was then averaged within each training paradigm along with non-CR trials and overlaid to investigate how MLI activity relates to differently timed CRs. Dividing the data with respect to CR amplitude involved aligning each trial by CS onset and sorting the behavioral data into equal subgroups with respect to CR amplitude above pre-CS baseline. Subgroup eyelid CRs and single unit data was then averaged and overlaid to investigate how MLI activity relates to CRs with different amplitudes. The absence of overlap in 95% confidence intervals between groups of average single unit activity indicated a significant difference.

#### Histology

After the conclusion of experiments the final tetrode placement was determined by making small marking lesions by passing 10 µA of anodal DC current for 10 s through tetrodes which yielded data. Animals were killed with an overdose of sodium pentobarbital and perfused intracardially with 0.9% saline (∼1.0 L) followed by 10% formalin (∼1.0 L). Heads were post fixed in formalin for at least 3 days after which tetrodes were removed and the brains were extracted. Brains were then cryo-protected in 30% sucrose in formalin for 3 days, embedded in an albumin gelatin mixture, and the cerebellum was sectioned using a freezing microtome at 40 µm. Tissue was mounted on slides and stained with cresyl violet, sections were then examined to determine the final location of each tetrode and this depth was compared with depth records from turning sessions to identify the location of unit recordings (**Supplementary Fig. 1a**).

### Computer simulations of the cerebellum

The details of the simulation used have been presented elsewhere^20–28^ and source code is available by contacting author MDM. The properties of the simulation are intended to emulate the synaptic organization and physiology of the cerebellum^1^ (Fig. 5a). As an approximation to the ratio of cell types within the cerebellum, the simulation implemented the following neuronal elements: 600 mossy fibers, 12000 granule cells, 900 Golgi cells, 96 basket cells (including 9 designated as PC-MLI), 240 stellate cells, 24 PCs, 8 deep cerebellar nuclei cells and four climbing fibers. These neurons were interconnected to emulate a parasagittal stripe, where all PCs receive input from climbing fibers of the same type – that is, those activated by an eyelid US and that control eyelid responses with their output. Each neuron was represented as a conductance-based calculation of membrane potential, with spikes determined by thresholds that varied according to recent spiking activity to implement the net effects of active conductances. Using 1 ms time steps, the change in membrane potential was calculated according to synaptic and leak conductances and to membrane capacitance. Spikes were initiated when membrane potential exceeded threshold. Threshold increased for each spike and returned exponentially to the baseline value. All synaptic conductances were based on time constants derived from *in vitro* studies, and the threshold properties of each representation was tuned to produce behavior that matches the *in vivo* characteristics of the target cell type.

The simulation represents the geometric relationships and divergence and convergence ratios of synaptic connections within the cerebellum by generating a two-dimensional array of granule cells, Golgi cells and mossy fiber glomeruli. For each type of connection an eligibility span was specified to represent the region of the array that the presynaptic neuron could potentially make contact with a post-synaptic target. These spans are based on published accounts of the spatial relationships of connections within the cerebellum. For example, since the axons of granule cells run transversely through the cerebellar cortex, the contact area for granule cells was a narrow rectangle. While this area constrained the range over which cells could make connections, the connections were determined randomly in a way that produced the known divergence and convergence ratios (for example, each granule cell could only receive four mossy fiber inputs).

Separate rules for plasticity were implemented at two types of synapses within the simulation: 1) the granule cell-to-PC synapses and 2) the mossy fiber-to-DCN synapses. A granule cell-to-PC synapse underwent LTD or LTP every time it fired a threshold burst of spikes, LTD occurred when this burst fell within a window between 300 ms and 100 ms prior to a climbing fiber input to the PC, otherwise LTP occurred. Mossy fiber-to-DCN synapses active within a time window of an abrupt pause in Purkinje cell activity underwent LTD whereas those active during strong Purkinje activity underwent LTP.

Eyelid conditioning was simulated by presenting to the simulation mossy fiber and climbing fiber inputs based on empirical recordings during eyelid conditioning. Each mossy fiber was assigned a background firing rate between 1 and 40 Hz. To mimic activation of mossy fibers as a conditioned stimulus, a randomly selected 3% of the mossy fibers were designated phasic CS mossy fibers and were active for a brief 100 ms burst at CS onset. Another randomly selected 3% were designated tonic CS mossy fibers and were activate at a rate between 80 and 100 Hz throughout the duration of the CS. All mossy fiber activity was stochastic with the target firing rate for any given time used to determine the probability of activating an excitatory conductance in ways that made the actual activity noisy. The activation of an excitatory conductance for the four climbing fibers served to mimic the presentation of the US. The averaged and smoothed activity of the eight deep nucleus neurons was used to represent the output of the simulation and the predicted “eyelid response” of the simulation.

To calculate the distribution of cross-correlations between PCs-basket cells activity in the simulation we did the following. We repeated 1000 times random draws of the same number of PC-BC pairs from the simulation, as we have total (64) in experimental data for ISIs 500, 700 and 750. These randomly drawn pairs were used to calculate cross-correlation values between PCs and basket cells average firing rates in the same fashion as with real data. For statistical comparison between correlation distributions of real data and simulations, we draw 1000 times from the simulation the same amount of PC-PC-MLI pairs as in real data and made a comparison using either paired Student’s t-test or two-sample Kolmogorov-Smirnov test for each draw.

## Data and code availability

Data that support the findings of this study and code used in this study are available from the corresponding authors upon reasonable request.

## ACKNOWLEDGMENTS

Financial support was provided by the Singapore Ministry of Education to G.J.A. (MOE2016- T2-1-097 and MOE2017-T3-1-002) and grants MH46904 and MH74006 to M.D.M.

## AUTHOR CONTRIBUTIONS

J.K. designed and did all cerebellar slice experiments, analyzed all data from these experiments, and wrote paper; G.J.A. helped design cerebellar slice experiments and wrote paper; H.E.H. designed and did all recording experiments during eyelid conditioning, analyzed data, and wrote paper; A.K. analyzed data and wrote paper; M.D.M helped design recording experiments during eyelid conditioning, performed simulations, analyzed data, and wrote paper.

## SUPPLEMENTARY INFORMATION

### Supplementary Text

During eyelid conditioning, a subset of PCs (eyelid PCs), which generate complex spikes in response to the unconditioned stimulus, show learned decreases in simple spike activity that highly correlate with eyelid conditioned responses (CRs) on a trial-by-trial basis^32^. Such learning-related plasticity could result from increases in the strength of the inhibitory synapse from MLIs to PCs (green in Fig. 5a) or from decreased excitation of PCs, due to weakened excitatory synapses onto PCs (blue in Fig. 5a). The latter would increase the activity of connected PC-MLIs receiving PC feedback inhibition and further inhibit neighboring PCs (as in Fig. 4d). To distinguish between these two hypotheses, the simulations compared different sites of learning-dependent synaptic plasticity^33, 34^: (1) parallel fiber (PF) inputs to PCs; (2) PF inputs to both PCs and MLIs; and (3) PF inputs to MLIs (Fig. 5a). These simulations exhibited CRs when learning-related plasticity occurred at PF-PC synapses (versions 1 and 2) but not when plasticity was restricted to PF-MLI synapses (version 3). Importantly, version (3) failed to support learning despite including feedback from the deep cerebellar nuclei, indicating that eyelid conditioning requires plasticity at PF-PC synapses. Although simulations can fail for myriad reasons, the learning inability displayed by simulations with plasticity only at MLIs suggests PC-to-MLI inhibitory feedback is more critical for motor learning. Therefore, during cerebellar learning, PC activity apparently decreases mainly because of reduced excitation at PF-PC synapses - due to long-term depression (LTD) - rather than from increased inhibition due to long-term potentiation (LTP) at PF-MLI synapses.

**Supplementary Figure 1.**
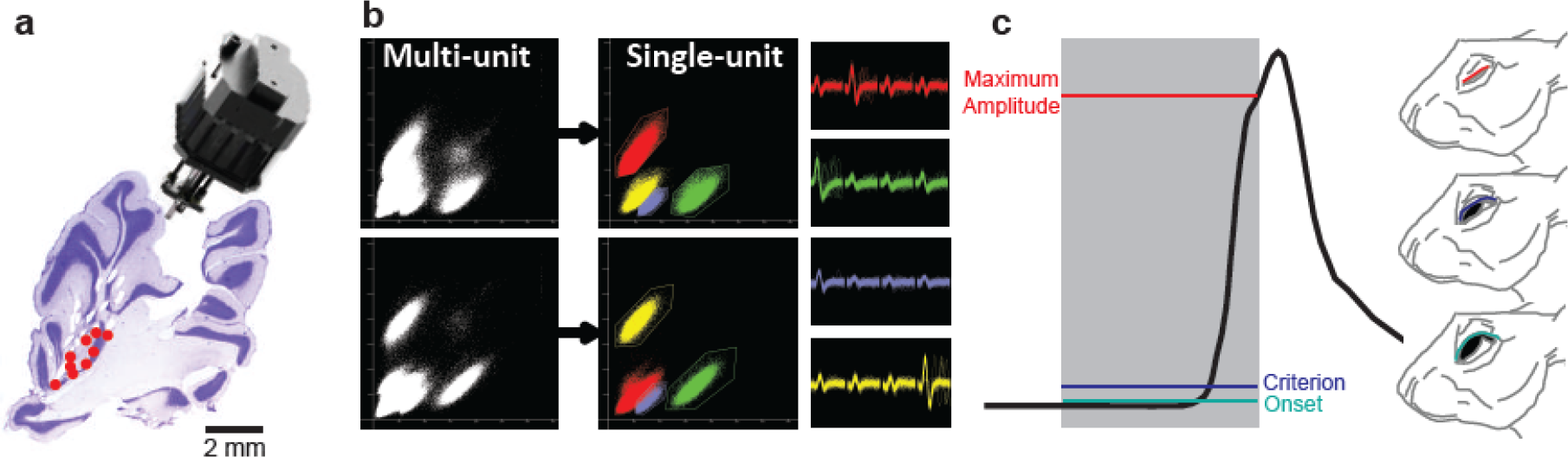
Single unit recordings in cerebellum and eyelid conditioning in the rabbit. (a) Sagittal histological section of the cerebellum showing tetrode tracks and the final location (red dots) of tetrodes that recorded eyelid MLIs and eyelid PCs. Our custom-made hyperdrive array is shown above the section. (b) Example of single units being isolated from a multi-unit recording via cluster cutting. The individual clusters and waveforms are color-coded to illustrate how unique features of the waveform across the four channels of the tetrode can be used to isolate single units from the multi-unit recording. (c)An exampleof a single conditioned eyelid response. Black line represents the position of the eyelid throughout the trial, while gray shading indicates the duration of the conditioned stimulus. The conditioned response is delayed relative to stimulus onset and peaks at the end of the ISI. Different response measurements are color-coded according to eyelid position (right).

**Supplementary Figure 2.**
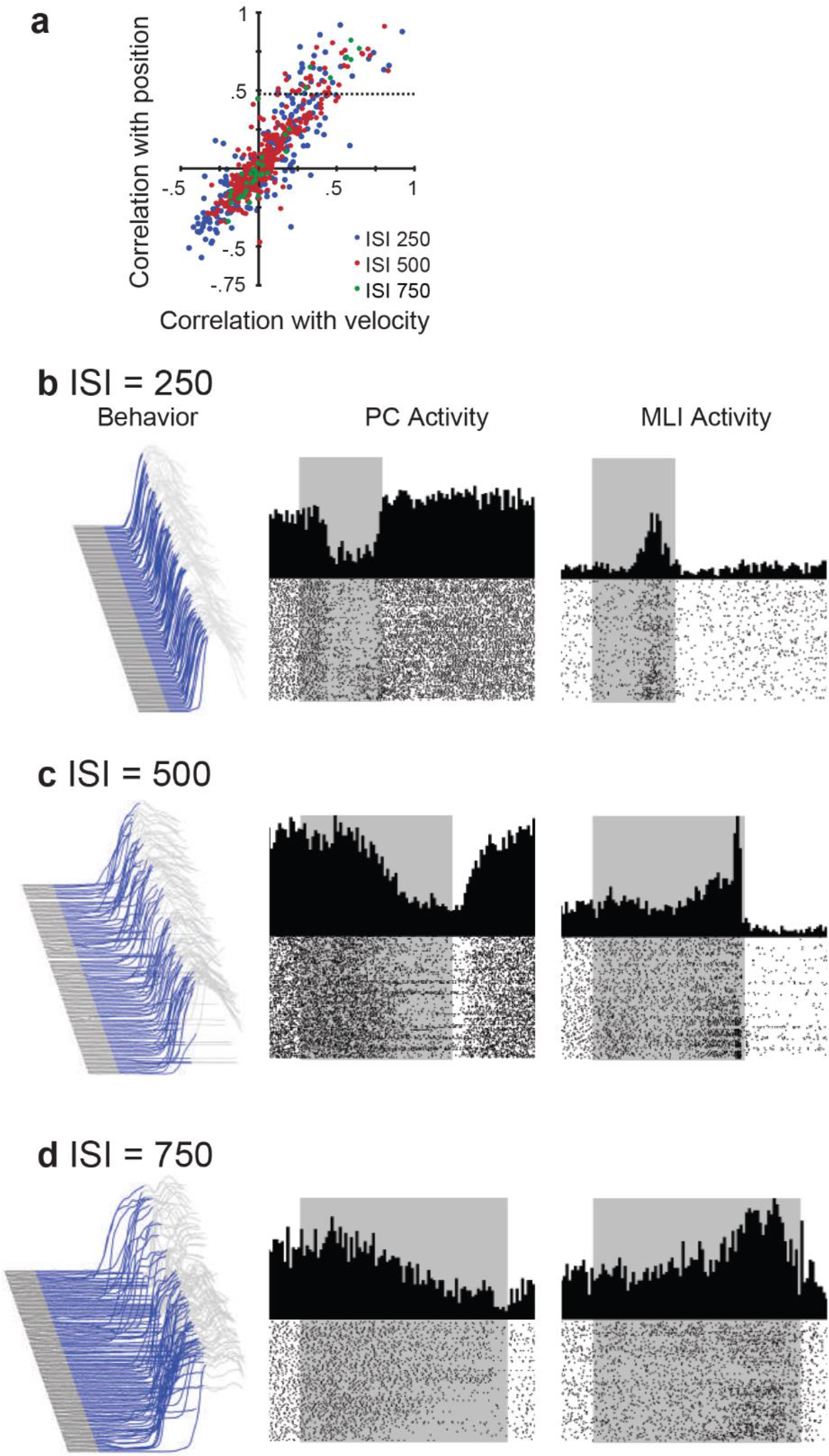
Relationship between simultaneously recorded activity of eyelid MLIs and eyelid PCs and conditioned eyelid responses. (a) Scatter plot showing the cross-correlation for all MLI recordings and either eyelid position or velocity at three different ISIs. Very few MLIs correlated more highly with velocity than position. All correlations above the dotted horizontal line were classified as putative eyelid PC-MLIs. (b-d} Eyelid movements and the activity of a PC and a putative PC-MLI simultaneously measured at different ISIs. Left - Eyelid movement during a conditioning session. Blue area indicates the duration of the tone stimulus (conditioned stimulus) and upward deflection is the conditioned response of the eyelid. Center - Peri-stimulus time histograms and raster plots of eyelid PC activity measured during the session shown at left. Right - Peri-stimulus time histograms and raster plots of putative eyelid PC-MLI activity measured during the same session. These are the same data shown in Figure 6 a-c and are shown here to illustrate how trial-to-trial variability in conditioned response timing can obscure the appearance of the tight relationship between eyelid MLIs and PCs; this relationship becomes clear once the data are sorted by time of onset of the conditioned response, as in Figure 6 a-c.

**Supplementary Figure 3.**
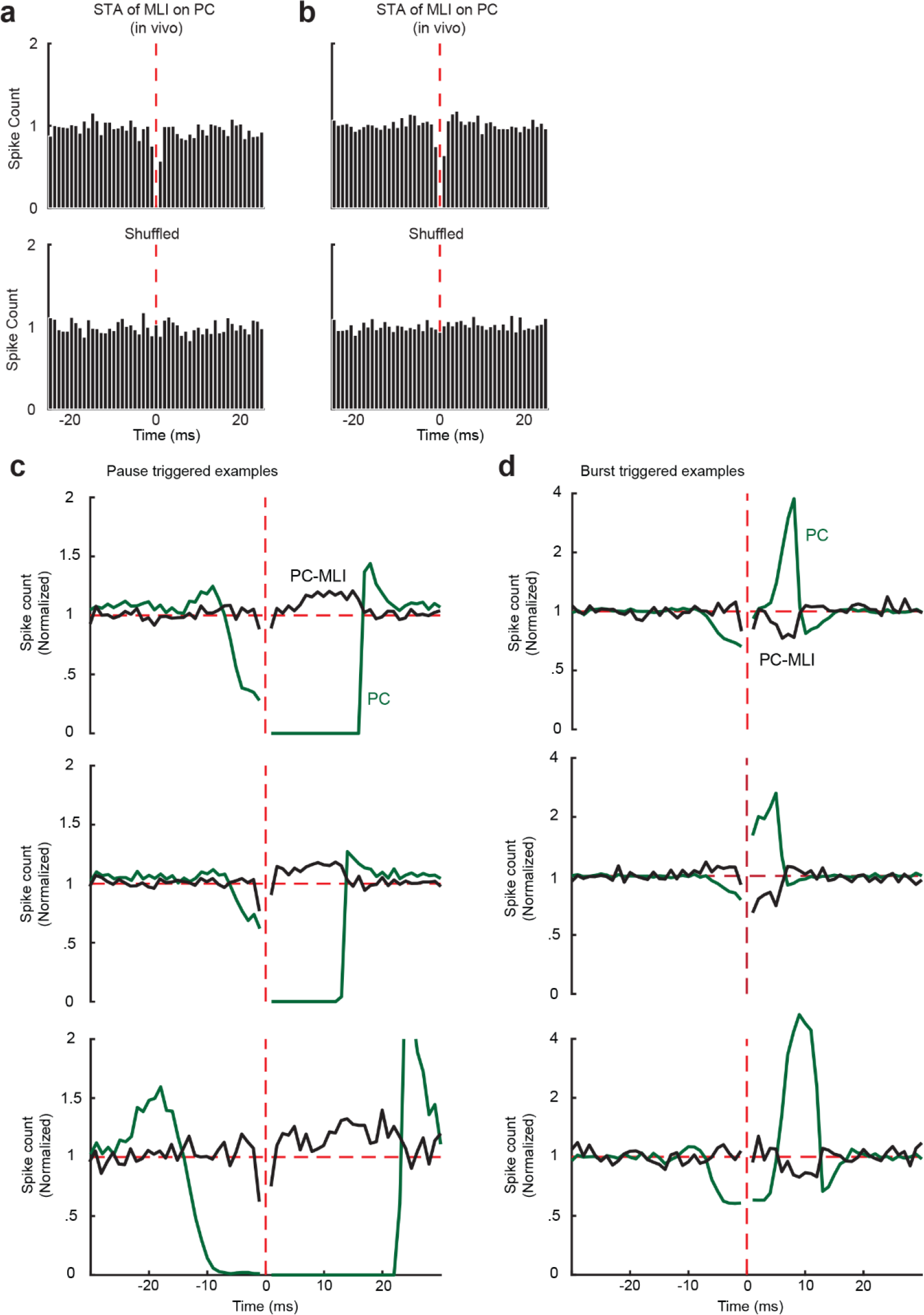
Relationship between baseline activity of PC-MLIs and PCs simultaneously recorded from the same tetrode. (a and b) Two examples of a spike-triggered average between MLI background activity triggered off PC simple spikes {vertical red line) recorded on the same tetrode. A total of six in vivo MLI/PC pairs showed a significant decrease In MLI activity after a PC spike. The post-spike decrease in MLI activity was abolished by shuffling the data. (c and d) Representative pause-triggered (c) and burst-triggered (d) cross-correlograms between background activity of putative PC-MLI (black) and eyelid PC simple spikes (green) recorded on the same tetrode for three different cell pairs. In all cases, pauses In PC simple spikes were associated with an Increase of PC-MLI activity, while bursts of PC simple spikes were associated with a decrease in PC-MLI activity.

**Supplementary Figure 4.**
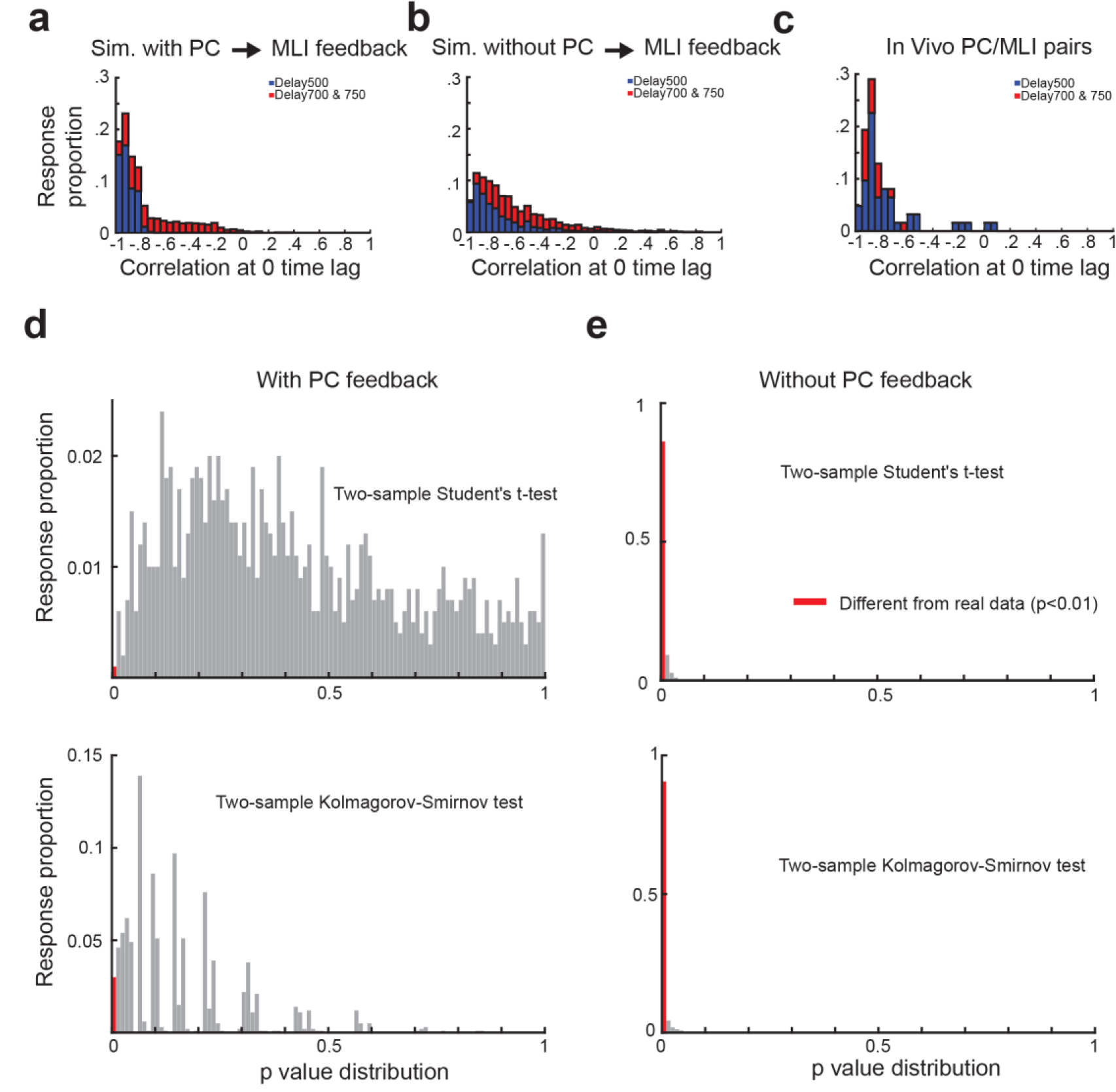
Statistical comparison of coupling between PC/PC-MLI activity In in vivo recordings and In simulations. (a-c) Relationship between simulated PC/PC-MLI pairs with and without PC feedback inhibition, as well as for in vivo pairs. Distributions of cross-correlations (at time lag = 0) between simultaneously recorded eyelid PC-MLI/PC pairs for IS Is 500, 700 and 750 ms for the computer simulation with PC to MLI collaterals (a), computer simulation without PC to MLI collaterals (b) and in vivo recordings (c).(d and e) Comparison of data from in vivo recordings in (c) and simulation with PC-to-PC-MLI feedback (a). We made 1000 draws from the simulation of 64 PC/PC-MLI pairs (equal to the number of pairs recorded in vivo). Each draw was compared with the distribution of PC/PC-MLI cross-correlations measured from in vivo data, (d) - Distribution of ρ values from all draws, using either two-sample Student’s t-test (top) or paired Kolmogorov-Smirnov test (bottom panel). Red bar indicates the fraction where there was a statistically significant difference (ρ<0.01) between real data and simulation, (e) - Same procedure applied to the simulation without PC-to-PC-MLI feedback (b). In this case, the vast majority of draws exhibited a significant difference In the correlation between activities of PC/PC-MLI pairs compared to the in vivo observations.

**Supplementary Figure 5.**
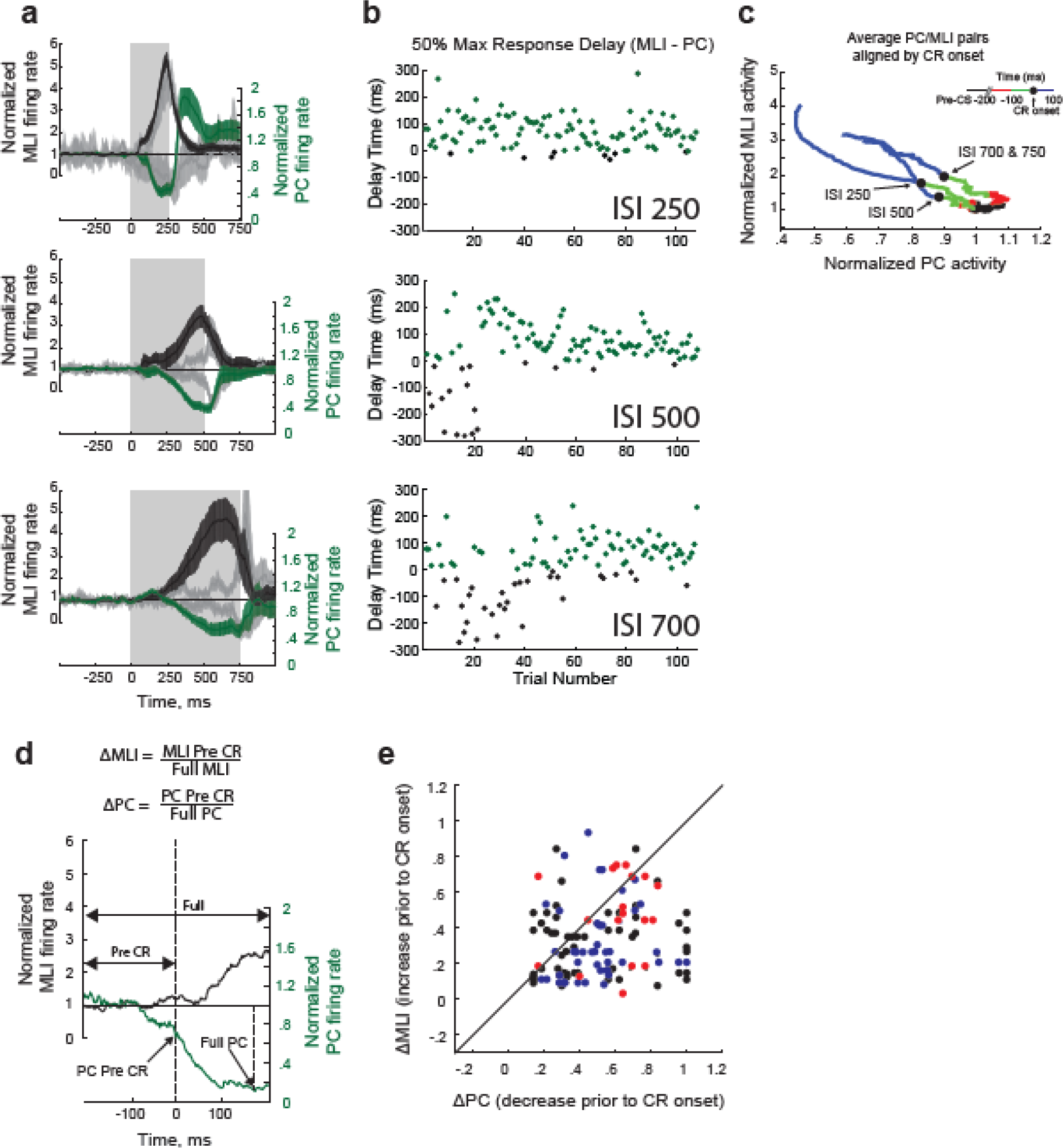
Relationship between averaged activity of pairs of eyelid PCs and putative PC-MLIs and quantification of this relationship for individual pairs. (a) Mean firing rate of all simultaneous recordings of PC/PC-MLI activity at three different ISIs (indicated by shaded rectangles). Mean PC-MLI activity (black lines) and PC activity (green lines) averaged across CR trials illustrates how the inverse activity relationship of the pairs shifts to match the training interval; shading represents 95% confidence intervals. Gray lines represent activity measured during non-response trials. (b) Trial by trial 50% maximum response delay between increases in PC-MLIs and decreases in simultaneously recorded eyelid PCs during eyelid conditioning at different ISIs. Negative numbers (black circles) indicate the PC-MLI reached 50% max response (increase) before the eyelid PC on that trial. Positive numbers (green circles) indicate the eyelid PC reached 50% response (decrease) before the PC-MLI. These 3 examples are from the data shown in Figure 7. (c) Color-coded plots for each ISI of the average normalized activity for all simultaneously recorded eyelid PC/PC-MLI pairs aligned to time of response onset (black dot). Green represents 100 ms before response onset and blue represents 100 ms after response onset in each case (inset). For each ISI, the trend was for PC activity to decrease before conditioned response onset and for the PC activity to continue to decrease while PC-MLI activity increased after response onset. (d) ΔPC and ΔMLI were calculated for each PC/PC-MLI pair whose activity was recorded simultaneously. For each pair, the change in PC activity (green) before response onset (PC pre CR) was divided by the change over the entire duration (full PC); the same analysis was done for each PC-MLI response (black). (e) A comparison of the pre-CR activity between eyelid PC/PC-MLI pairs. The ΔPC to ΔMLI ratio was plotted for each simultaneously recorded pair where points along the diagonal line indicate equal change before conditioned response onset. Deviations below the diagonal line indicate that the eyelid PC changed more than the simultaneously recorded activity of the putative PC-MLI, while deviations above the diagonal line indicate that the putative PC-MLI changed more than the eyelid PC before response onset. Each paired recording is color-coded according to the ISI (black = ISI 250, blue = ISI 500, red = ISI 700 & 750).

**Supplementary Figure 6.**
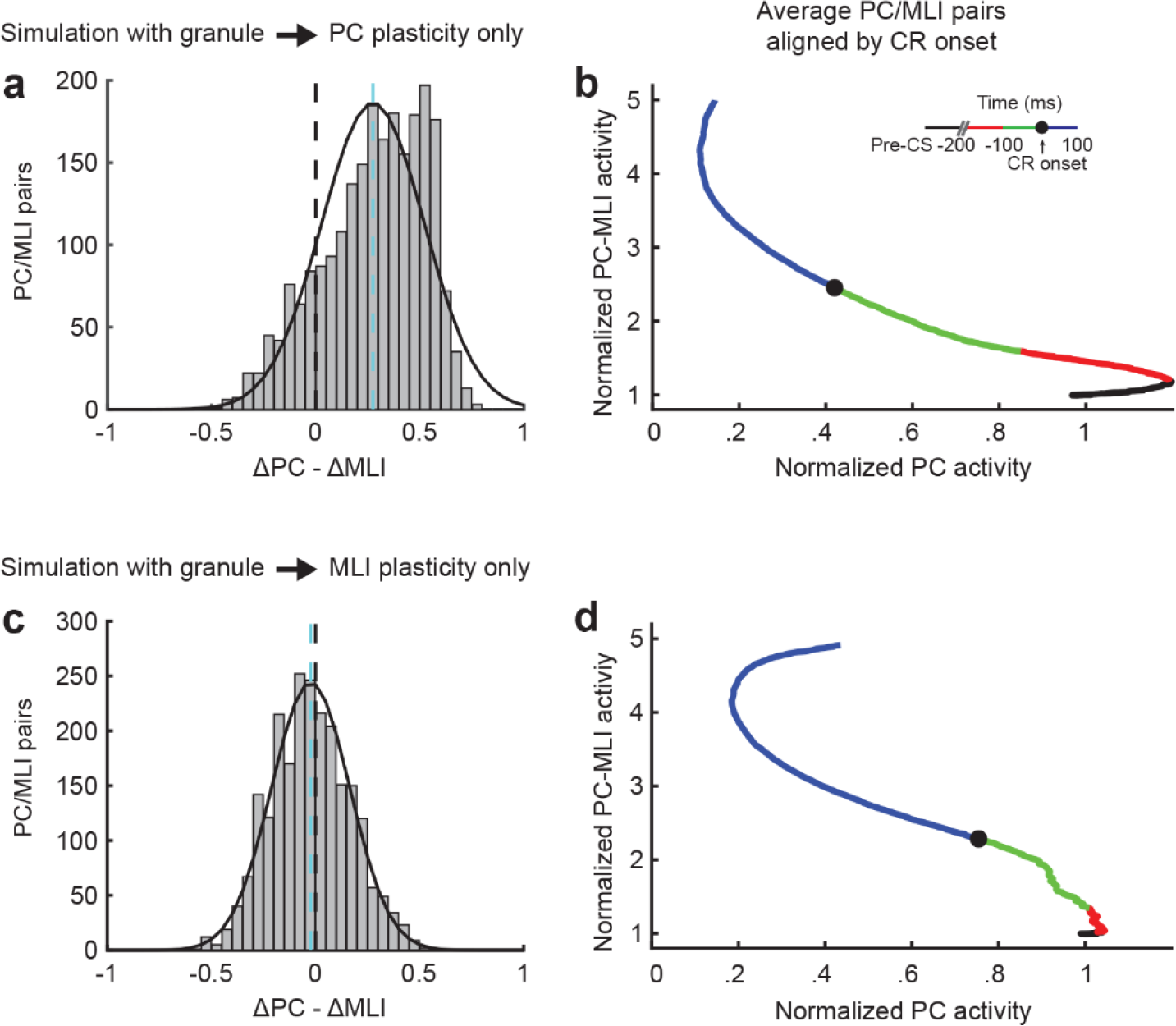
Relationship between activity of PC and PC-MLI pairs from simulations featuring different sites of circuit plasticity. (a) Distribution of the relative latency of eyelid PCs decreases in activity relative to increases in PC-MLIs (ΔPC -Δ MLI) in simulations with plasticity only at granule cell-to-PC synapses. The distribution ofΔPC -ΔMLI differences (mean indicated by cyan dashed line) is shifted to the right of zero (black dashed line), indicating that PC activity usually decreases more than the PC-MLI activity increases relative to conditioned response onset. (b) Color-coded plots of the time of average normalized activity for all eyelid PC/PC-MLI pairs from the simulation with plasticity only at granule cell-to-PC synapses, aligned to response onset (black dot). Green represents 100 ms before response onset and blue represents 100 ms after response onset in each case (inset). PC activity tended to decrease before the time of CR onset and then continued to decrease after CR onset, while the PC-MLI activity increased later. This was similar to the trend observed in vivo (Supplementary Figure 6b). (c) Same distribution as in (a), from a simulation with plasticity only at granule cell-to-MLI synapses. (d) same analysis as in (b), for the results of simulation with plasticity only at granule cell-to-MLI synapses. The trend was for the PC-MLI to increase its activity before CR onset and the PC to then decrease its activity after CR onset.

